# Targeting mTORC2 in lung squamous cell carcinoma improves anti-tumor immunity through the PSGL-1-VISTA axis

**DOI:** 10.1101/2025.04.23.650229

**Authors:** Verra M. Ngwa, Yoonha Hwang, Wenqiang Song, Deanna N. Edwards, Jin Chen

## Abstract

Targeted therapies have improved survival for lung adenocarcinoma patients. However, similar advances are lacking for lung squamous carcinoma (LUSC). Advances in immunotherapy have shown some promise, but the overall response rate remains low in LUSC. Here, we demonstrate that the mTORC2 signaling pathway represents an actionable target in LUSC to improve anti-tumor immune responses. We show that genetic alterations affecting the mTORC2 pathway are common among patients with LUSC tumors, and targeting mTORC2 reduces LUSC tumor growth in mouse models. Transcriptomics reveal that mTORC2-deficient LUSC cells exhibit reduced expression of glycolytic and hypoxia-related genes. In agreement, loss of mTORC2 signaling decreases lactate levels in tumor-interstitial fluid, creating reduced acidity within the tumor microenvironment. Interestingly, mTORC2-deficient LUSC cells also exhibited reduced expression of the pH-sensitive VISTA ligand PSGL-1 in a HIF-2α dependent mechanism. LUSC patients, but not those with LUAD, display a positive correlation in expression between HIF-2α and PSGL-1, suggesting a distinct association among mTORC2, HIF-2α, and immune responses in LUSC. Indeed, mTORC2 loss-of-function enhanced CD8^+^ T cell activation in tumors, while use of anti-VISTA immunotherapy reduced LUSC tumor burden only in the presence of intact mTORC2 signaling. Collectively, these data describe an important role of mTORC2 signaling in LUSC tumors and demonstrate the therapeutic potential of targeting the mTORC2/PSGL-1/VISTA axis in patients that are non-responsive to current therapies.

## Introduction

Lung squamous cell carcinoma (LUSC), which makes up about 30% of non-small cell lung cancer (NSCLC), represents a highly metastatic subclass of lung cancer that is strongly associated with smoking (1). Despite the success of targeted therapies in NSCLC lung adenocarcinoma, similar advances have not successfully materialized in LUSC. The high mutational burden in NSCLC favors the generation of neoantigens that can be recognized by cytotoxic T cells, leading to successful integration of immune checkpoint blockades (ICBs) that are approved as first-line treatment for patients with high PD-L1 expression and second-line agents for patients who fail chemotherapy (2–4). Despite these advances, the response rates to immunomodulatory therapies remain less than 50% in the first line with high PD-L1 and 20-30% in the second line (2,5)

Intensive efforts have been made in recent years to improve immunotherapy outcomes. While much attention remains on PD-1 and PD-L1, recent advances have identified expression of the checkpoint molecule VISTA in most lung cancers, including LUSC (6). VISTA acts as a checkpoint receptor, selectively binding to the P-selectin glycoprotein ligand-1 (PSGL-1) at an acidic pH (7). VISTA and PD**-**1 play non-redundant roles in regulating T cell responses, and VISTA is recognized as a potential mediator of resistance to anti-PD**-**1 and anti-CTLA4 immune therapies in prostate cancer patients (8). However, the functional consequence of VISTA in LUSC has not been investigated.

Characterization of LUSC patient tumors in large cohort studies has previously identified the PI3K-mTOR-Akt pathway as a significantly affected pathway in LUSC (2,4). The mammalian target of rapamycin (mTOR) is a serine-threonine kinase that exists in two distinct complexes, mTORC1 and mTORC2 (9). In addition to mTOR, the mTORC2 complex is composed of Rictor, MLST8, and SIN1 among others, while mTORC1 consists of mTOR, Raptor, MLST8, and PRAS40. Although MLST8 is present in both complexes, it plays a critical role in the stability and function of mTORC2 but not mTORC1 (10–12). The importance of mTORC1 has been extensively studied in cancer models, but the impact of mTORC2 is poorly understood (13,14). In LUSC, PTEN deletion is common, which results in hyperactivation of mTORC2 even in the absence of growth factor stimulation (4,15). Downstream activation of mTORC2 leads to phosphorylation and activation of the AGC kinases AKT, SGK, and PKC, which function to promote cell survival, cytoskeletal organization, and cellular metabolism. Additionally, PI3K-mTORC2 signaling has been implicated in regulating the expression of HIF2α, one of the master regulators of cellular responses in solid tumors (16)Therefore, the mTORC2 pathway may play a central role in LUSC tumorigenesis.

In this report, we demonstrate that elevated mTORC2 signaling is associated with worse outcomes in LUSC. In mouse models, targeting the mTORC2 pathway reduces LUSC tumor growth and improves markers of T cell activation. We also show that mTORC2 loss within LUSC tumor cells reduces glycolysis and decreases interstitial lactate to create a less acidic tumor microenvironment. Transcriptomics confirmed reduced expression of glycolytic genes after mTORC2 loss and revealed a decrease in *SELPLG*/PSGL-1 expression that is dependent on *EPAS1*/HIF2α. Blocking the PSGL-1/VISTA interaction using an anti-VISTA antibody significantly inhibited LUSC tumor growth in wild-type but not mTORC2-deficient tumors, suggesting that targeting the mTORC2-PSGL-1-VISTA pathway may offer a new therapeutic avenue for LUSC.

## Methods

### Cell Lines and Cell Culture

HCC2814 cells were obtained from the Hamon Cancer Center Collection (University of Texas Southwestern Medical Center, Dallas, TX). H596 and KLN205 were purchased from ATCC. JH716 cells were provided by Chad Pecot (University of North Carolina, Chapel Hill, NC). All tumor cells were maintained in Dulbecco’s modified Eagle’s medium (DMEM) (Corning #10-013-CV) supplemented with 10% fetal bovine serum (FBS), 100 U/ml penicillin, and streptomycin at 37 °C in a humidified atmosphere containing 5% CO_2_ and 95% air. *Mycoplasma* testing was performed on cultured cells every 6 months using the MycoStrip-*Mycoplasma* Detection kit (rep-mys-10, Invitrogen).

### Generation of CRISPR- KO cell lines

Human and mouse knockout cell lines were generated as previously described (10). In brief, sgRNAs targeting *MLST8* were designed, amplified, and cloned into lentiCRISPR v2. Using standard protocols, lentiviruses were packaged in HEK293FT cells by transfecting cells with CRISPR or expression plasmids together with psPAX2 (lentiviral packaging) and pCMV-VSV-G (envelope) plasmids at 1:1:1 molar ratio using the Lipofectamine 2000 Reagent. Media was changed after 6 hrs of transfection and virus was collected after 48∼72 hrs. Indicated cells were infected with virus in the presence of 8 μg/ml Polybrene for 24 hrs and selected with 1-2 μg/ml Puromycin or 20-40 μg/ml Blasticidin until non-transduced control cells died before being used for experiments.

The *Pten* and *Mlst8* knockout KLN205 cell line was generated using the transfection of Cas9 ribonuleoprotein (RNP) complex technique. Cas9 protein (Integrated DNA Technologies, Inc.) was assembled with sgRNA for *Pten* or *Mlst8* at 1:1 molar ratio in OptiMEM media. The Cas9/sgRNA RNP complex was reverse transfected into KLN205 cells with RNAiMAX (Thermofisher scientific). After 48 hrs post-transfection, transfected cells were seeded into a 96-well plate by limited dilution. Once cloned cells were expanded, double knockout cells (KLN205 Pten^null^ Mlst8^null^) were screened by western blot for PTEN and MLST8. sgRNA sequence were AATTAATGGGTGCGTTCACC for *Mlst8* and AATTAATGGGTGCGTTCACC for *Pten*. Wild type Mlst8 or empty vector was re-expressed in the KLN205 Pten^null^ Mlst8^null^ double knockout cell line to generate KLN205 Pten^null^ Mlst8^null^ WT or Mlst8-KO.

To overexpress full-length ovalbumin (OVA), JH716 cells were transduced with lentivirus from pLVX-Hygro-IRES-OVA as previously described (17). Single clones were obtained through limiting dilutions. JH716-OVA Mlst8 loss of function mutant cells (JH716-OVA-Mlst8-KO) and WT (JH716-OVA-WT) were generated as described above.

### Mouse Model

All mice were housed in a non-barrier animal facility. Rag1-deficient (C57BL/6, Rag1^null^) mice were obtained from Jackson Laboratories, and were maintained in sterile housing. DBA/2 mice were purchased from Envigo or The Jackson Laboratories. OT-I (C57BL/6) transgenic mice were purchased from The Jackson Laboratories.

### Tumor models and treatment regimens

KLN205 Pten^null^ Mlst8^null^ WT or Mlst8-KO knockout cells (5 x10^5^) or H596 WT or MLST8-KO cells (5 x 10^6^) were implanted subcutaneously into the dorsal flanks of recipient mice (DBA/2 or Rag1^null^, respectively). For HIF2α rescued experiment, 5 x10^5^ KLN205-Pten^null^ Mlst8-KO expressing empty vector (Mlst8-KO + EV) or constitutive active HIF2α cDNA (Mlst8-KO + HIF2α_CA) (Addgene) were implanted into DBA/2 mice subcutaneously (SQ) or via tail vein administration. All SQ tumors were measured using calipers, and tumor volume was calculated using the formula V= 0.5* (L x W^2^).

For drug studies, KLN205-Pten^null^ WT or Mlst8-KO (5 x10^5^) were seeded into the lung via tail vein injection and the mice were randomized and treated with IgG control (10mg/kg, BioXCell #BE0091; Lebanon, NH, USA) or anti-VISTA (10mg/kg, BioXCell, #BE0310; Lebanon, NH, USA) starting on day 4. The mice were treated every 3-4 days for 7 doses via intraperitoneal injection. Animals were euthanized on day 25, and their lungs were flushed at least twice with PBS via the left ventricle of the heart. The weight of the lungs was measured, and detailed images were captured using an Olympus stereo microscope.

### Flow Cytometry

Flow cytometry experiment was performed as previously described (18). Briefly, tumors were dissociated in RPMI-1640 media (Corning #MT10040CV, Corning, NY, USA) supplemented with 5% FBS, 1 mg/ml collagenase IA (Sigma-Aldrich #C9891, St Louis, MO, USA), and 0.25 mg/ml DNase I (Sigma-Aldrich #DN25) for 30 minutes at 37°C followed by filtering the digested tissue through a 70-µm strainer. Red blood cells were lysed using ACK Lysis Buffer (KD Medical #RGF-3015, Columbia, MD, USA) and samples washed with PBS before staining with Ghost Dye Violet V510 (Tonbo Biosciences #13-0870, San Diego, CA USA) to exclude dead cells. After washing with buffer (0.5% BSA, 2mM EDTA in PBS), samples were blocked in anti-CD16/32 mouse Fc block (Tonbo Biosciences #70-0161) and cell surface proteins were analyzed using antibodies against: CD45, TCRβ, CD4, CD8a, CD25, CD162, PD-1, and/or CD107a. Intracellular staining for GZMB was accomplished using a Cytofix/Cytoperm solution kit (BD 554714, Franklin Lakes, NJ, USA) on paraformaldehyde-fixed cells, according to the manufacturer’s protocol. Fluorescence minus one (FMO) or IgG controls were included in the gating controls when needed using splenocytes or tumor cell suspensions. Flow cytometry data was obtained on a BD Fortessa FACS Diva software and analyzed using FlowJo software (v10). Antibodies used in flow panels are detailed in Supplemental Table 1.

### RNA Sequencing

RNA sequencing was performed as previously described (18). In brief, RNA was extracted from HCC2814 sgControl (WT) or sgMLST8 (MLST8-KO) using the Direct-zol RNA Miniprep Kit (Zymo Research, Irvine, CA, USA) according to manufacturer’s instructions. RNA sequencing was performed by BGI Americas (Cambridge, MA, USA) using the DNBSEQ platform. Raw data was filtered after sequencing to remove reads with high rates of unknown bases, low quality reads, and reads of adapter sequences. Clean reads were aligned to the reference genome (Homo Sapiens, version GCF_000001405.38_GRCh38.p12) using HISAT and aligned to reference genes using BowTie2. Using the Dr. Tom platform (BGI), differentially expressed genes (DEG) were identified using DESeq2 (q value < 0.05). Pathway enrichment analysis was performed against Molecular Signatures Database (MSigDB) Hallmark gene sets using ShinyGO Software (0.77) (19).

### Gene alteration analysis

110 input gene identifiers from the human REACTOME_PI3K_AKT_SIGNALING_IN_CANCER gene set obtained from MSigDB (20–22) were used to examine their alterations from 5 TCGA Pan-cancer data sets [**NSCLC**: TCGA PanCancer Atlas (n=1144); **LUSC:** CPTAC (n=80), TCGA Firehose Legacy (n= 511); **LUAD**: CPTAC (n=110), TCGA Firehose Legacy (n=586)] using tools developed by cBioPortal (https://www.cbioportal.org/). The percentage of total alteration frequency was calculated from one or more datasets, as indicated. Genes in the mTORC2 pathway were further evaluated on cBioPortal to illustrate the genetic alterations in the PI3K-mTORC2-AKT pathway.

For survival analyses, relapse-free survival (RFS) and overall survival (OS) based on mean mTORC2-related gene (*PIK3CA*, *PTEN* (inverted), *AKT1*, *AKT2*, *AKT3* and *RICTOR*) expression in stage II-IV lung squamous cell carcinoma were downloaded from KM plot (http://www.kmplot.com) (23). Automatic cutoff was used to define “high” versus “low” expression from RNA-seq data, with cutoff values defined as RFS=1422.17 (range = 536-11116) and OS = 1793.67 (range = 536-11116). Similar survival analyses were performed using 24 mTORC1 gene identifiers obtained from the human REACTOME_MTORC1_MEDIATED_ SIGNALLING and seven mTORC1-related gene (*RPTOR, RPS6, RPS6KA1, AKT1S1, ELF4EBP1, ELF4EBP2, ELF4EBP3*) expression in stage II-IV lung squamous cell carcinoma were downloaded from KM plot. Same as above, automatic cutoff was used to define “high” versus “low” expression from RNA-seq data, with cutoff values defined as RFS=4068.73 (range = 2774 – 11354) and OS = 4679.68 (range = 2774 - 11354).

### Western Blot and Co-immunoprecipitation (Co-IP)

Western blot and co-IP assays were performed as previously described (10). Cultured tumor cells were lysed with CHAPS lysis buffer (40 mM Tris, pH 7.5, 120 mM NaCl, 1 mM EDTA, 0.3% CHAPS) and RIPA buffer supplemented with protease inhibitors and phosphatase inhibitors (Complete Mini and PhosStop inhibitor cocktail, Roche). Protein concentration of each sample was determined by Pierce BCA Protein Assay kit. Equal amounts of protein extracts were mixed with 4x Laemmli sample buffer, separated by electrophoresis on a 10% or a gradient 4%–20% SDS-PAGE gel, and transferred onto nitrocellulose membranes. Membranes were blocked with 5% skim milk in TBS/T buffer and incubated with corresponding primary antibodies and IRdye-conjugated secondary antibodies. Immunoreactivity was detected using the Odyssey scanner (Li-cor Biosciences, Lincoln, NE, USA). To perform immunoprecipitation, equal amounts of input lysates (2-4 mg) were incubated with the primary antibodies (1-2 μg) for 2 hrs to overnight at 4°C. Protein G Dynabeads were added, and lysates were incubated for 1 hr and washed four times with CHAPS lysis buffer.

### Immunohistochemistry

Immunohistochemical staining of tumor lung tissue was performed on formalin-fixed, paraffin-embedded blocks as previously described(18). Briefly, serial sections were deparaffinized in xylene and rehydrated through a graded series of ethanol concentrations. Antigen retrieval was achieved by microwave-boiling the sections for 10 min in citrate buffer. Sections were allowed to cool down for 30 minutes followed by PBS washes. Intrinsic peroxidase activity was blocked with 3% hydrogen peroxide in methanol for 10 min. Sections were blocked using goat serum (2.5%; Sigma) solution, and the optimally diluted primary antibodies (CD8a, 1:100, # 14-0808-82, eBioscience, San Diego, CA, USA; Cleaved Caspase-3, 1:100, #9664L, CST, Danvers, MA, USA) were applied to cover the specimen and incubated at 4°C overnight. After three washes in 1x PBS for 5 min each, slides were incubated with biotinylated secondary antibody at room temperature for 1hr. Following three additional washes in PBS, slides were incubated with streptavidin peroxidase reagent # SA-5704-100 (Vector Laboratories, Newark, CA, USA) for 45 min followed by additional PBS washes. Sections were developed with DAB substrate (#550524, BD Pharmingen) solution per manufacturer’s protocol and counter stain lightly with hematoxylin. Slides were dehydrated and mount with Cytoseal (#48212-196, VWR, Radnor, PA, USA). Images of at least five fields of view were obtained using an Olympus inverted fluorescence microscope (40x). The intensity of the staining as well as the percentage of positive cells was recorded.

### Seahorse analysis

Cellular and mitochondrial bioenergetics of human and murine LUSCs were measured using the Seahorse XFe/XF-24 Extracellular Flux Analyzer (Seahorse Bioscience, North Billerica, MA, USA). Briefly, human (HCC2814 WT or MLST8-KO cells; H596 WT or MLST8-KO) and murine (KLN205-PTEN^null^ WT or Mlst8-KO) cells were plated in XF-24 cell culture plates (1 × 10^5^ cells/well) in 100 μL DMEM medium as mentioned above for 1hr before adding 400µL complete medium and incubated overnight. Glycolysis stress and Mito stress tests were performed as per the manufacturer’s instructions (103020-100 and 103015-100, Agilent Technologies). For glycolysis stress test, tumor cells were washed in XF Assay media supplemented with 2 mM glutamine and 1 mM pyruvate. Glycolytic stress test drugs were prepared in prepared XF media at the following final concentrations: glucose (10 mM), oligomycin (0.25 uM), and 2-Deoxy-D-glucose (2-DG) (10 mM) To perform the mitochondrial stress test, tumor cells were washed in XF Assay Media supplemented with 10 mM glucose, 2 mM glutamine and 1 mM pyruvate. Mitochondrial stress test drugs were prepared in XF media at the following final concentrations: Oligomycin (1.5 μM), Carbonyl cyanide-4 (trifluoromethoxy) phenylhydrazone [(FCCP), 2.0 μM], and Rotenone/Antimycin A [(Rot/AA), 0.5 μM].

### Quantitative real-time polymerase chain reaction

Total RNA was isolated and reversely transcribed using the RNeasy kit (Qiagen, #74104, Hilden, Germany) and iScript cDNA synthesis kit (Bio-Rad, # 1708891, Hercules, CA, USA) according to the manufacturers’ directions. qPCR was performed using the SYBR Green PCR Master Mix (#4368706, Thermo Fisher Scientific, Walthman, MA USA) on StepOne system (Applied Biosystems, Foster City, CA, USA). Each condition was assayed in triplicate. The primer sequences used for each gene are presented on Supplemental Table 2. Quantitation was performed using the ΔΔ*C*_t_ method.

### Tumor interstitial fluid metabolic assays

Tumor interstitial fluid was collected from tumors as previously described (17) with some modifications. Tumors were harvested, weighed, and then washed with PBS to remove excess blood from the tumor surface. Excess PBS was carefully removed by blotting. Tumors were placed in a 20 μm pluriStrainer (pluriSelect, #43-10020-40, Leipzig, Germany) mounted in a microcentrifuge tube, then centrifuged at 100g at 4°C for 30 minutes. The cleared interstitium was deproteinated and lactate was detected using the Lactate Assay Kit (Sigma-Aldrich, MAK064) according to the manufacturer’s instructions. Known lactate standards were used to calculate concentrations.

### Cytotoxicity assay

In vitro cytotoxicity assay was described previously (17) with modifications. Briefly, splenocytes from DBA/2 mice were activated with anti-CD3 (2.5μg/mL) and anti-CD28 (2.5μg/mL) for 48hrs. CD8^+^ T cells were isolated using mouse CD8a microbeads (Miltenyi Biotec, 130-045-201) according to the manufacturer’s instructions. Conditioned media was prepared by culturing 4x10^5^ KLN205-Pten^null^ WT or Mlst8-KO cells in a 12-well plate for 48 hrs. For the co-culture, 6x10^4^ KLN205-Pten^null^ WT or Mlst8-KO cells were plated into a 24-well plate and allowed to attach overnight. Tumor cells were stained with CellTracker Green (CTG, 2.5μM; #C7025, Thermo Fisher Scientific) for 30 min and washed prior to adding T cells. Tumor cells were co-cultured in the presence or absence of activated CD8^+^ T cells (6x10^5^) in the appropriate conditioned medium for an additional 48 hrs. Cells were collected using accutase and stained with 7-AAD (1μg/mL, Life Technologies #A1310) for 30 min at 4°C. Dead (7-AAD+) CellTracker Green-positive tumor cells were determined by flow cytometry.

Splenocytes were harvested from OT-1 mice and activated with SIINFEKL peptide (1μg/mL; Invivogen #vac-sin) for 48 hrs. 2x10^5^ JH716-OVA-sgControl or sgMlst8-KO cells were cultured in a 12-well plate for 48 hrs to collect conditioned media. For the co-culture, 1x10^4^ JH716-OVA-sgControl or sgMlst8-KO cells were allowed to attach in a 96-well plate overnight. CD8+ T cells purified as described above were co-cultured with tumor cells at a 1:1 ratio in the appropriate conditioned media. CyQuant LDH cytotoxicity assay (Invitrogen, #C20300) was performed in triplicate as described by the manufacturer, and cytotoxicity was calculated, accounting for non-T cell mediated tumor cell death. To evaluate cytotoxicity in response to VISTA or PSGL-1 neutralization, 2x10^5^ JH716-OVA-sgControl cells were co-cultured with activated CD8+ OT-1 T cells as described above in the presence of anti-VISTA (100ug/mL), anti-PSGL-1 (100ug/mL), or an IgG control (100ug/mL) for 48 hrs.

### Statistics

GraphPad Prism software (10.0.3) was used to generate graphs and performed statistical analyses. Summary data are expressed as the mean ± SEM. For in vivo tumor experiments, data are reported from 2-3 independent experiments, where each dot represents a mouse. Statistical comparisons between two groups were performed using a nonparametric, unpaired Student’s t-test. For multiple comparisons, one-or two-way analysis of variance (ANOVA) was performed, and individual comparisons were evaluated using either Tukey’s, Dunnett’s, or Sidak’s post hoc analysis as indicated. Outliers were excluded using the ROUT method (Q=5%). Differences with a p-value less than 0.05 were considered statistically significant. For Kaplan-Meier survival curve analysis, log-rank analysis was performed between groups using the Mantel-Cox method. Hazard ratios (HR) and 95% confidence intervals (CI) were determined using the log-rank test. Pearson correlation (r) was performed on TCGA datasets.

## Study approval

Animal care and experimental procedures were performed under protocols approved by Vanderbilt University’s Institutional Animal Care and Utilization Committee (protocol number: IACUC M2200094).

## Data Availability

RNA-sequencing data has been deposited in the Gene Expression Omnibus (GEO) and are available under the accession numbers GSE269711. Further inquiries can be directed to the corresponding author.

## Results

### Genetic alterations in PI3K-mTORC2-AKT pathway correlate with patient survival in human lung squamous cell carcinoma

To identify genetic vulnerability of the mTORC2 pathway in LUSC, we first assess the frequency of molecular alterations in PI3K and AKT signaling in the two predominant non-small cell lung cancer (NSCLC) subtypes, lung squamous cell carcinoma (LUSC) and lung adenocarcinoma (LUAD). Analysis of multiple LUSC and LUAD datasets reveals that the PI3K-AKT pathway is highly amplified in LUSC compared to LUAD (Fig. 1A). More specifically, most LUSC patients carry one or more genetic alterations in the mTORC2-related genes (Fig. 1B). Of these alterations, amplification of *PIK3CA*, *RICTOR*, *AKT1*, *AKT2* or *AKT3* were the most frequent followed by deep deletion of *PTEN* (Fig. 1B), all likely representing hyperactivation of mTORC2 signaling.

**Figure 1:**
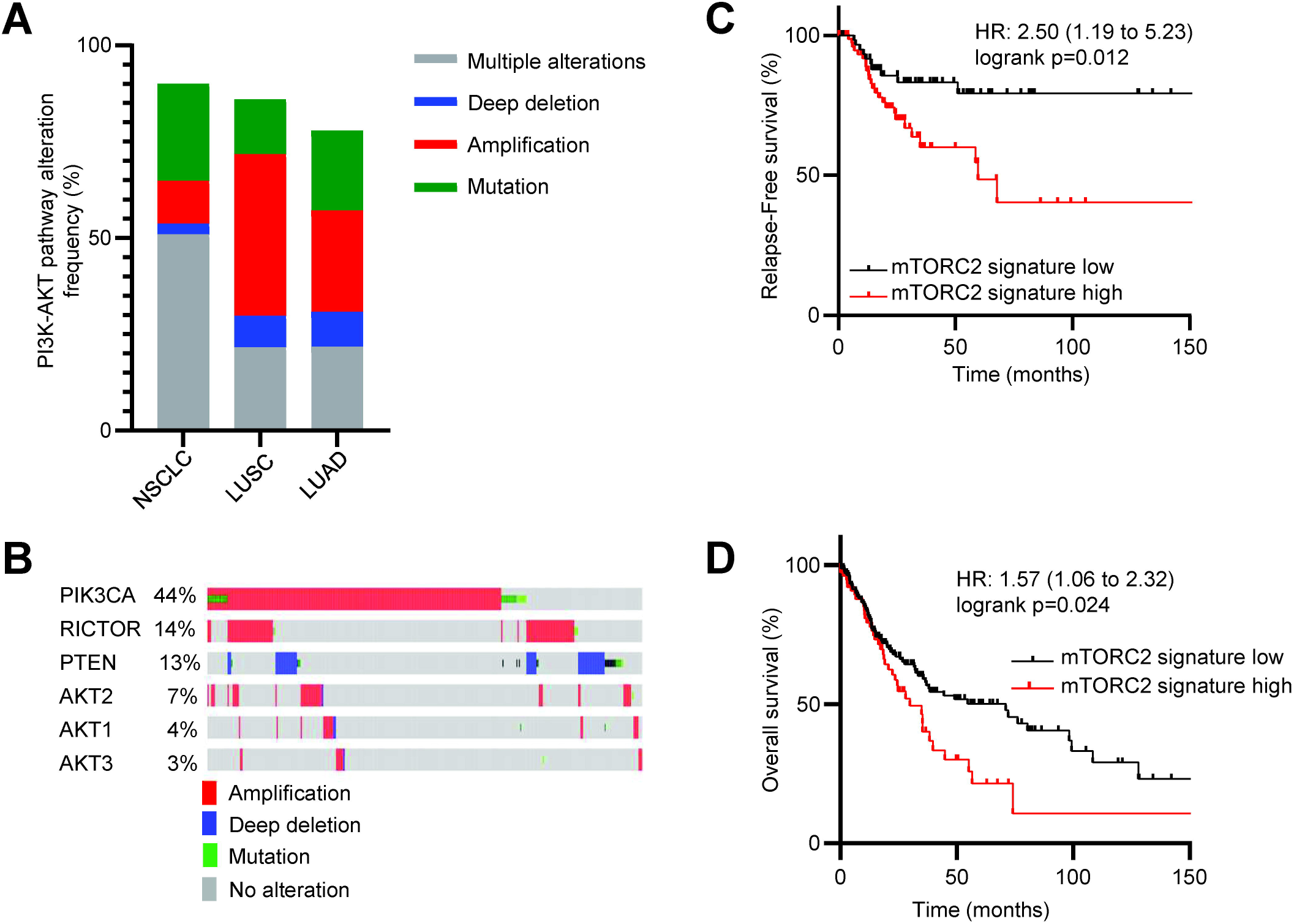
Genetic alterations of the PI3K- mTOR- AKT pathway in squamous cell lung cancer. (**A**) Molecular alteration in non-small cell lung cancer (NSCLC) using the human gene set for REACTOME_PI3K_AKT_SIGNALING_IN_CANCER pathway from 5 cancer datasets [**NSCLC**: TCGA PanCancer Atlas (n=1144); **LUSC:** CPTAC (n=80), TCGA Firehose Legacy (n= 511); **LUAD**: CPTAC (n=110), TCGA Firehose Legacy (n=586)] using tools developed by cBioPortal. (**B**) Analysis of mTORC2 related genes (mTORC2 signature: *PIK3CA*, *RICTOR*, *PTEN*, *AKT1-3*) that are amplified, deleted, and mutated from the LUSC studies in panel **A** above [**LUSC:** CPTAC (n=80); TCGA Firehose Legacy (n= 511)]. (**C-D**) Kaplan-Meier (KM) of stage II-IV plot displaying the probability of relapse-free survival (**C**) and overall survival (**D**) with association of the mTORC2 related genes illustrated in **B** using lung squamous cell carcinoma RNA-seq dataset (n=141, relapse-free survival and n= 249, overall survival) downloaded from kmplot.com.

Given these genetic alterations may affect expression levels, we next assessed the functional consequence of mTORC2-related genes (*PIK3CA*, *RICTOR*, *PTEN*, *AKT1-3*, collectively referred to as mTORC2 signature) expression on LUSC survival. We found that high expression of mTORC2 signature genes correlates with unfavorable survival probabilities in advanced LUSC patients, an important finding given that most LUSC patients are not diagnosed until more progressive disease has developed (Fig. 1C, D). To ensure that these changes were mostly driven by mTORC2 pathway in these LUSC patients, we analyzed Reactome genes involved in mTORC1 signaling as well as specific downstream genes of the mTORC1 pathway. In either case, we found that mTORC1 signaling was not associated with survival in LUSC patients, suggesting that LUSC is highly driven by the mTORC2 pathway (Fig. S1).

### mLST8 loss selectively inhibits mTORC2 signaling and tumor growth in vivo

Although MLST8 is a common component for both mTORC1 and mTORC2 (24), structural studies revealed that MLST8 is more integrated in the mTORC2 complex (25–27). Indeed, we and others showed that MLST8 loss selectively disrupts mTORC2 integrity, signaling and kinase activity in LUAD and early development models, recapitulating loss of the unique mTORC2 component, Rictor (10,12). Therefore, we deleted MLST8 (MLST8-KO) in human LUSC cell lines to test the effect of mTORC2 loss on LUSC. Importantly, we utilized human HCC2814 and H596 cells that harbor *PTEN* deletion and *PIK3CA* mutation, respectively, representing models of genetic mTORC2 hyperactivation in human tumors. Loss of MLST8 disrupts association of mTOR kinase with RICTOR but not RAPTOR, consistent with MLST8’s selective role in mTORC2 signaling in LUAD (10). Additionally, MLST8 loss significantly reduced phosphorylation of AKT at S473, a direct mTORC2 substrate site, but does not affect phosphorylation of the mTORC1 substrate S6K1 (Fig. 2A). To model human disease using a syngeneic murine line, we generated a *Pten* knockout in the murine LUSC cell line KLN205 (KLN205-Pten^null^), Consistent with the results in human LUSC cell lines, KLN205-Pten^null^ WT, but not KLN205-Pten^null^ Mlst8-KO, retained mTOR kinase interactions with RICTOR, as well as phosphorylation of AKT at S473 (Fig. 2B) but not affecting mTORC1 targets pS6K, pS6 or p4EBP, either in complete growth media or in serum-starved and restimulated conditions (Fig. S2A), confirming the selectivity of MLST8 for mTORC2 in LUSC.

**Figure 2:**
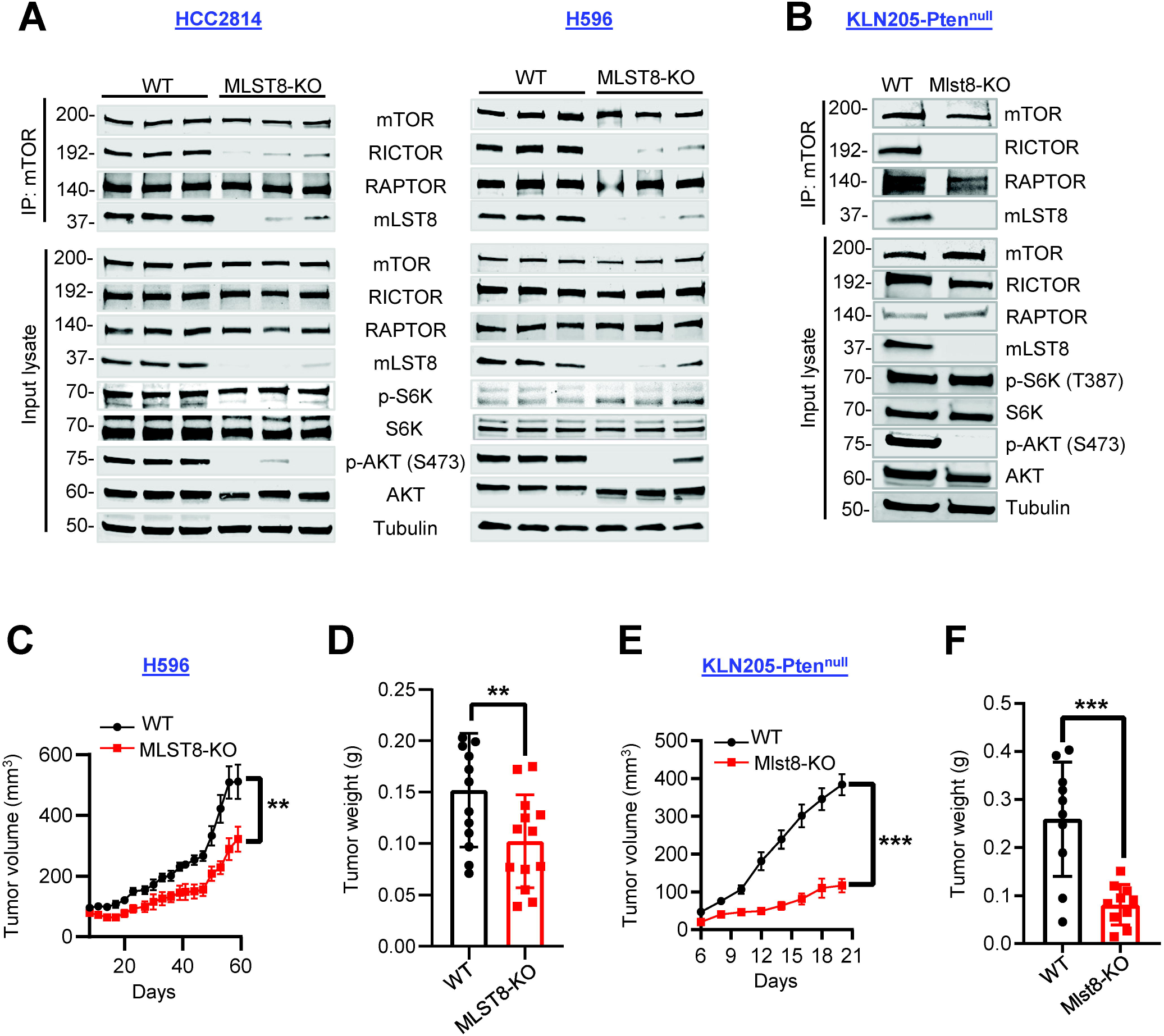
Selectively targeting mTORC2 inhibits LUSC tumor growth in vivo. (**A**) mTOR immunoprecipitate (IP) or cell lysates from human HCC2814 (PTEN null) and H596 (PIK3CA E545K) [WT (sgControl #1, #2, #3) and KO (sgMLST8 #1, #2, #3)] and (**B**) mouse KLN205-Pten^null^ WT or Mlst8-KO cells were assessed by western blot. (**C-D**) 5x10^6^ H596 WT or MLST8-KO cells were implanted subcutaneously in Rag1-deficient (Rag1^null^) mice. Tumor growth was monitored, and tumor volume calculated (**C**). Tumor weight was measured at harvest (n=12-13 mice per group) (**D)**. (**E**-**F**) 5x10^5^ KLN205-Pten^null^ WT or Mlst8-KO cells were implanted in DBA/2 mice. Tumor growth curves (**E**) and tumor weight (**F)** were measured as in **C** and **D** (n=10 mice per group). All data are presented as mean ± SEM from two or three independent experiments. Each dot represents a mouse. p-values were determined by 2-way ANOVA with Sidak multiple comparisons post hoc (**C** and **E**) or two-tailed unpaired Student *t* test (**D** and **F**). **p<0.01, ***p<0.001.

To determine the role of mTORC2 in LUSC in vivo tumor growth, we injected WT or MLST8-KO tumor cells into the flanks of recipient mice. Human HCC2814 and H596 tumors were evaluated in immune-deficient Rag1^null^ hosts that lack mature lymphocytes to create a permissive environment for xenograft growth (28). HCC2814 cells did not grow in vivo, but MLST8-KO cells did show a modest decrease in cell viability in vitro (Fig. S2B). In H596 cells, loss of MLST8 reduced tumor growth that was also associated with reduced cell viability in vitro (Figs. 2C, D; S2C). More strikingly, Mlst8 knockout of KLN205-PTEN^null^ cells markedly decreased tumor growth in immune-competent syngeneic DBA/2 mice, despite it having no effect on cell viability in vitro (Figs. 2E, F; S2D, E). These results suggest that loss of mTORC2 function in LUSC cells decreases tumor growth.

### Loss of mTORC2 in tumor cells enhances T cell effector function in vivo

To dissect the potential role of tumor cell mTORC2 on anti-tumor immune responses, we first used adoptive T cell transfer to introduce an adaptive immune system in human LUSC xenografts. WT or MLST8-KO H596 cells were implanted into the flank of Rag1^null^ recipients, and pre-activated CD8^+^ T cells isolated from C57BL/6 OT-1 mice were adoptively transferred once the tumors were established (Fig. 3A). Tumors were harvested, and the immune cell populations were analyzed by flow cytometry in tumor single cell suspensions. Although overall CD45^+^ immune cells and infiltrated CD8^+^ T cells were unchanged, CD8^+^ T cells in MLST8-KO tumors had increased expression of the activation marker CD25^+^ and the degranulation marker CD107a^+^ compared to WT controls (Figs. 3B-D, S3A), suggesting that reduced mTORC2 activity in tumor cells may create a more favorable tumor microenvironment for CD8^+^ T cell function.

**Figure 3:**
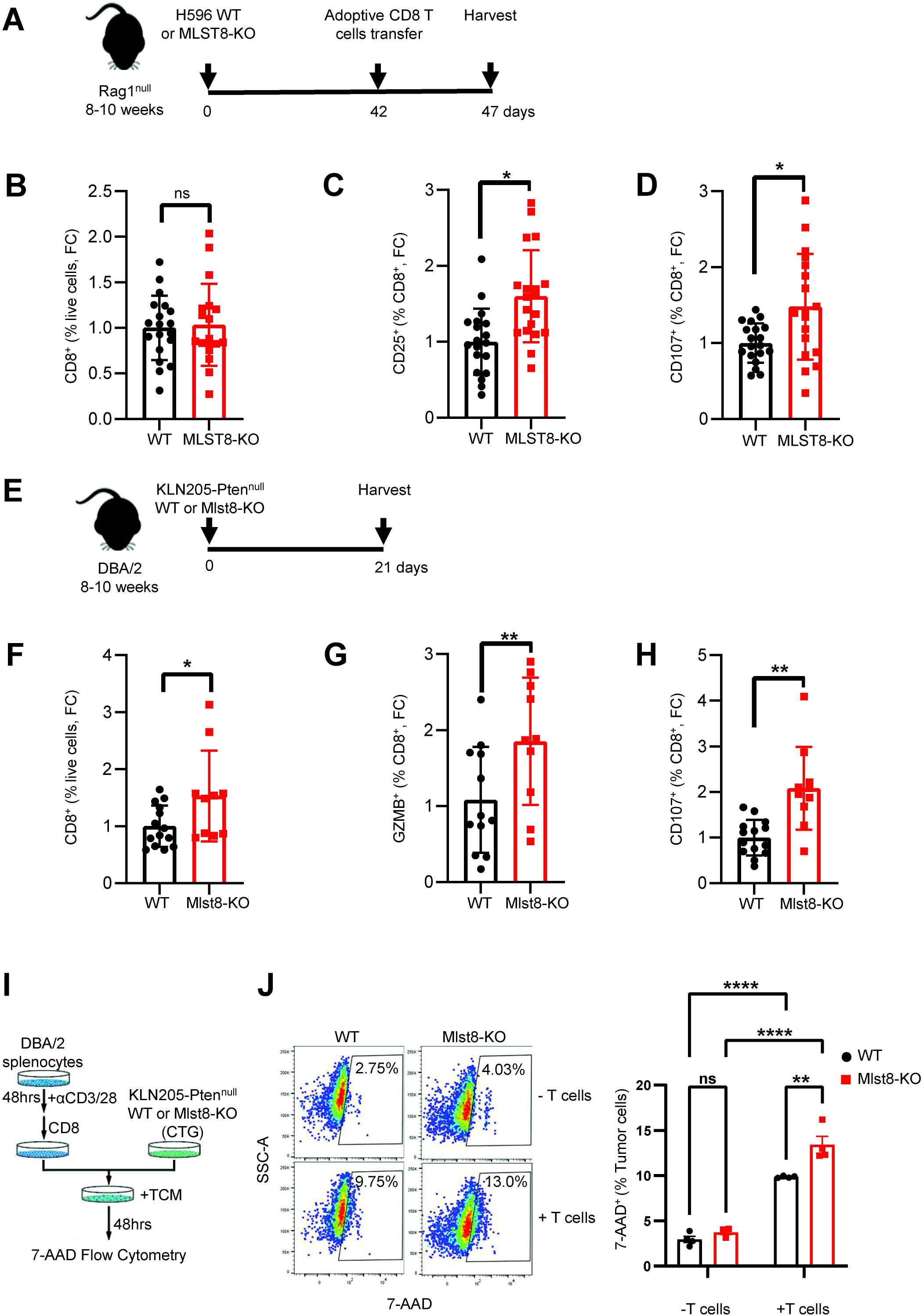
MLST8/mTORC2 loss improves effector function of cytotoxic T cells. (**A**) Schematic diagram of experimental design showing H596 tumor allograft in Rag1^null^ mice. (**B**) Flow cytometric analysis of CD8^+^ T cells in WT and MLST8-KO tumors. (**C-D**) Flow cytometric analysis of CD25^+^ (**C**) and CD107a^+^ (**D**) in CD8^+^ T cells of WT and mLST8-KO tumors. (**E**) Schematic diagram of experimental design showing KLN205-Pten^null^ WT or Mlst8-KO tumor allograft in DBA/2 mice. (**F-H**) Flow cytometric analysis of CD8^+^ in live cells (**F**), GZMB^+^ (**G**), and CD107a^+^ (**H**) in CD8^+^ T cells of WT and Mlst8-KO tumors. Each dot represents a mouse. (**I-J**) Cytotoxicity was determined using a co-culture assay. (**I**) Schematic of co-culture is shown. (**J**) CellTracker Green (CTG)-stained KLN205-Pten^null^ WT or Mlst8-KO cells were co-cultured in the presence or absence of CD8^+^ T cells in tumor conditioned medium (TCM) (n=3). All data are presented as mean ± SEM from two or three independent experiments. p values were determined by two-tailed unpaired Student *t* test. *p<0.05, **p<0.01, **p<0.001, ns: not statistically significant, FC: Fold change

Next, we evaluated how tumor cell mTORC2 activity impacts T cell responses in a fully immunocompetent murine model. KLN205-Pten^null^ WT and Mlst8-KO cells were implanted as indicated in Figure 3E, and flow cytometry was performed on harvested tumors. Immune analysis of harvested tumors revealed no significant changes in the numbers of CD45^+^, B, or NK cells (Fig. S3B-D) in Mlst8-KO tumors, while CD8^+^ and CD4^+^ T cells were increased (Figs. 3F, S3E). We further observed a significant increase in granzyme B (GZMB^+^) and CD107a^+^ CD8^+^ T cells, indicating increased cytolytic activity of CD8^+^ T cells in Mlst8-KO tumors (Fig. 3G-H). Indeed, in a direct co-culture of tumor cells with T cells, we observed an increase in tumor cell killing on Mlst8-KO tumor cells, compared with WT controls (Fig. 3I-J). In agreement with increased T cell activation, a greater percentage of CD4^+^ and CD8^+^ T cells from Mlst8-KO tumors expressed the immune checkpoint receptor PD-1 (Fig. S3F-G), whereas VISTA^+^ CD8^+^ or CD4^+^ T cells were moderately decreased in the Mlst8-KO tumors compared to WT controls (Fig. S3H-I). We further utilized the murine LUSC cell line JH716 (29) overexpressing the full-length ovalbumin protein (JH716-OVA) and co-cultured with activated OT-I CD8+ T cells isolated from OT-I transgenic mice (Fig. S3J). Again, there was a significantly increased tumor cell killing with Mlst8-KO compared to control (Fig. S3K). Collectively, these data indicate that Mlst8/mTORC2 loss in tumor cells enhances CD8^+^ T cells effector function and antitumor immune response.

### mTORC2 loss reduces tumor cell glycolysis and extracellular lactate

Tumor cell metabolism is a significant modulator of anti-tumor immune responses within the tumor microenvironment (17,30) and mTORC2 is known to regulate cellular metabolism (31,32). Recent studies have shown that LUSC cells have a higher glucose flux rate than LUAD (33,34), suggesting that mTORC2 hyperactivation may drive glycolysis in LUSC. Indeed, we observed that the phenol red pH indicator in the culture medium of MLST8-KO HCC2814 cells appears less yellow, or less acidic, than WT control cells at the same cell density (Fig. S4A). Lactate production from glycolytic reprogramming in cancer cells is a major contributor to extracellular acidification (35). Therefore, we assessed the extracellular acidification rate (ECAR) in MLST8-deficient HCC2814 cells as a readout of glycolytic function. Relative to control, overall ECAR is significantly decreased in MLST8-KO cells, suggesting that loss of MLST8/mTORC2 signaling reduces glycolysis in LUSC cells. Additionally, reductions in glycolytic capacity were observed, indicating that MLST8-deficient cells are less able to utilize glucose (Fig. 4A, B). Consistent with lower extracellular acidification, lactate concentration in the culture medium of MLST8-KO cells was significantly reduced (Fig. 4C). Similar reductions in glycolytic ECAR and extracellular lactate are observed in MLST8-deficient H596 and KLN205-Pten^null^ Mlst8-KO cells (Fig. 4D-H), while lactate levels were decreased in tumor interstitial fluid of KLN205-Pten^null^ Mlst8-KO tumors (Fig. 4I). Interestingly, HCC2814 MLST8-KO cells exhibit a moderate reduction in oxygen consumption rate, suggesting that cellular respiration may be somewhat impacted by targeting mTORC2 (Fig. S4B, C). Therefore, these results suggest that targeting mTORC2 may reduce tumor cell glycolysis and lower extracellular levels of lactate, creating a less acidic tumor microenvironment.

**Figure 4:**
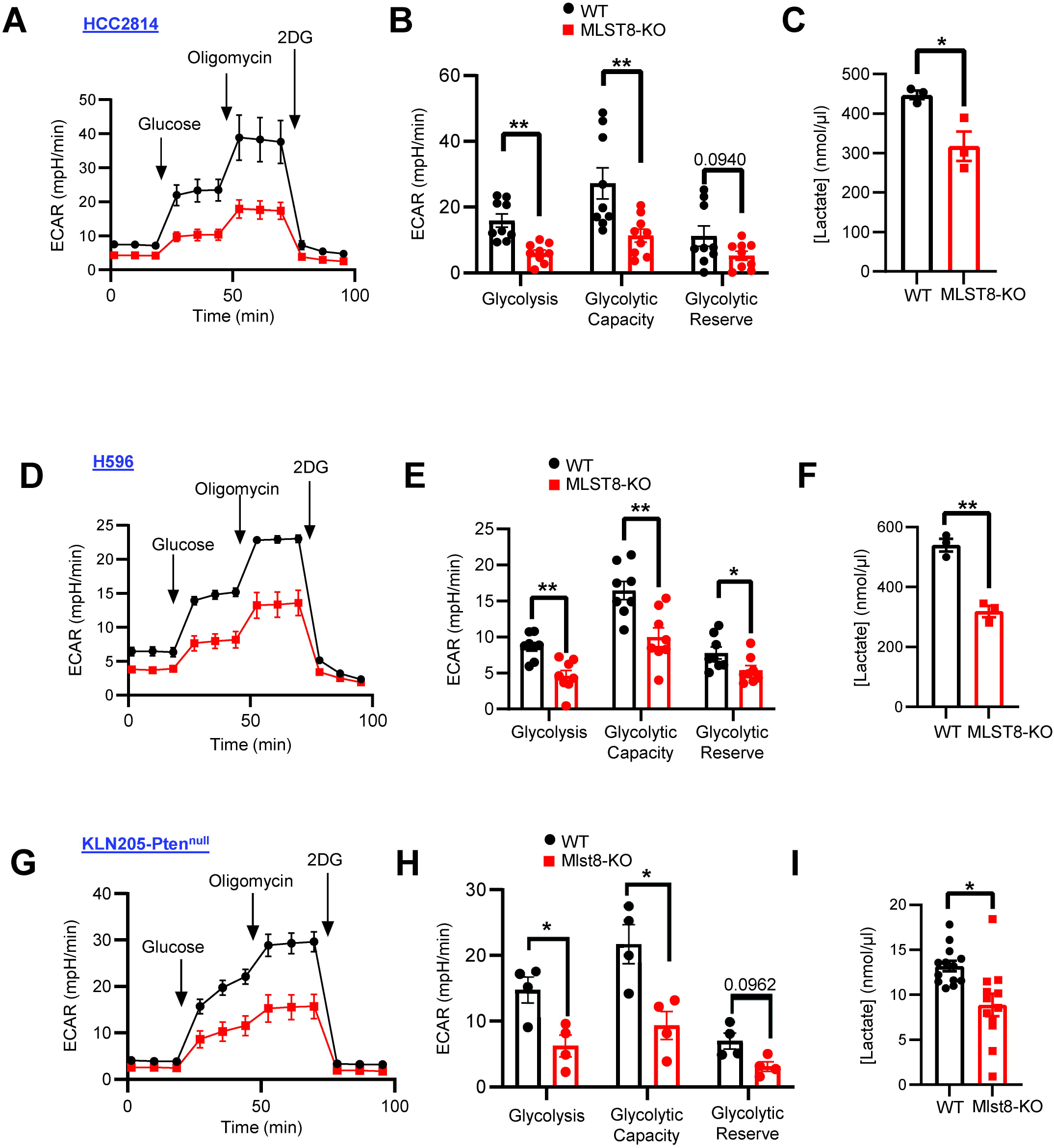
Targeting mTORC2 inhibits glycolysis and lactate secretion. (A-C) Seahorse analysis of (A) extracellular acidification rate (ECAR), (B) glycolysis, glycolytic capacity, and glycolytic reserve and (C) in vitro lactate concentrations in HCC2814 WT vs MLST8-KO. (D-F) Seahorse analysis of (D) extracellular acidification rate (ECAR), (E) glycolysis, glycolytic capacity, and glycolytic reserve and (F) in vitro lactate concentrations in H596 WT vs MLST8-KO. (G-I) Seahorse analysis of (G) extracellular acidification rate (ECAR), (H) glycolysis, glycolytic capacity, and glycolytic reserve and (I) *in vivo* tumor interstitial lactate concentrations in KLN205-Pten^null^ WT vs Mlst8-KO. All data are presented as mean ± SEM from two or three independent experiments. p values were determined by two-tailed unpaired Student *t* test. *p<0.05, **p<0.01.

### mTORC2 regulates PSGL-1 expression via HIF2α

To dissect mechanisms by which mTORC2 regulates tumor metabolism and immune regulation, we performed RNA-seq on HCC2814 WT and MLST8-KO cells. Pathway analysis revealed enrichment of downregulated differentially expressed genes (DEGs) involved in glycolysis in MLST8-KO cells (Fig. 5A, B). Indeed, qRT-PCR confirmed decreases in expression of the glycolytic genes *HK2, GLUT1*, and *LDHA* (Fig. 5C), consistent with a glycolytic defect in MLST8-KO cells. Strikingly, we also discovered that MLST8-deficient cells exhibit a significant decrease in expression of the *SELPLG* gene that encodes PSGL-1, a VISTA ligand at acidic pH (7) by RNAseq, qRT-PCR (Fig. 5A, C) and flow cytometry (Fig. 5D). However, we did not observe significant changes in SIGLEC-5, *VISR*/VISTA, or *CD279*/PD-L1 in MLST8-KO tumor cells, compared to WT controls (Fig. S5A-D). Given that the PSGL-1/VISTA interaction only occurs at an acidic pH (7), these data suggest that decreases in glycolytic genes and PSGL-1 may work in concert to improve anti-tumor immune responses upon targeting mTORC2 in LUSC tumors.

**Figure 5:**
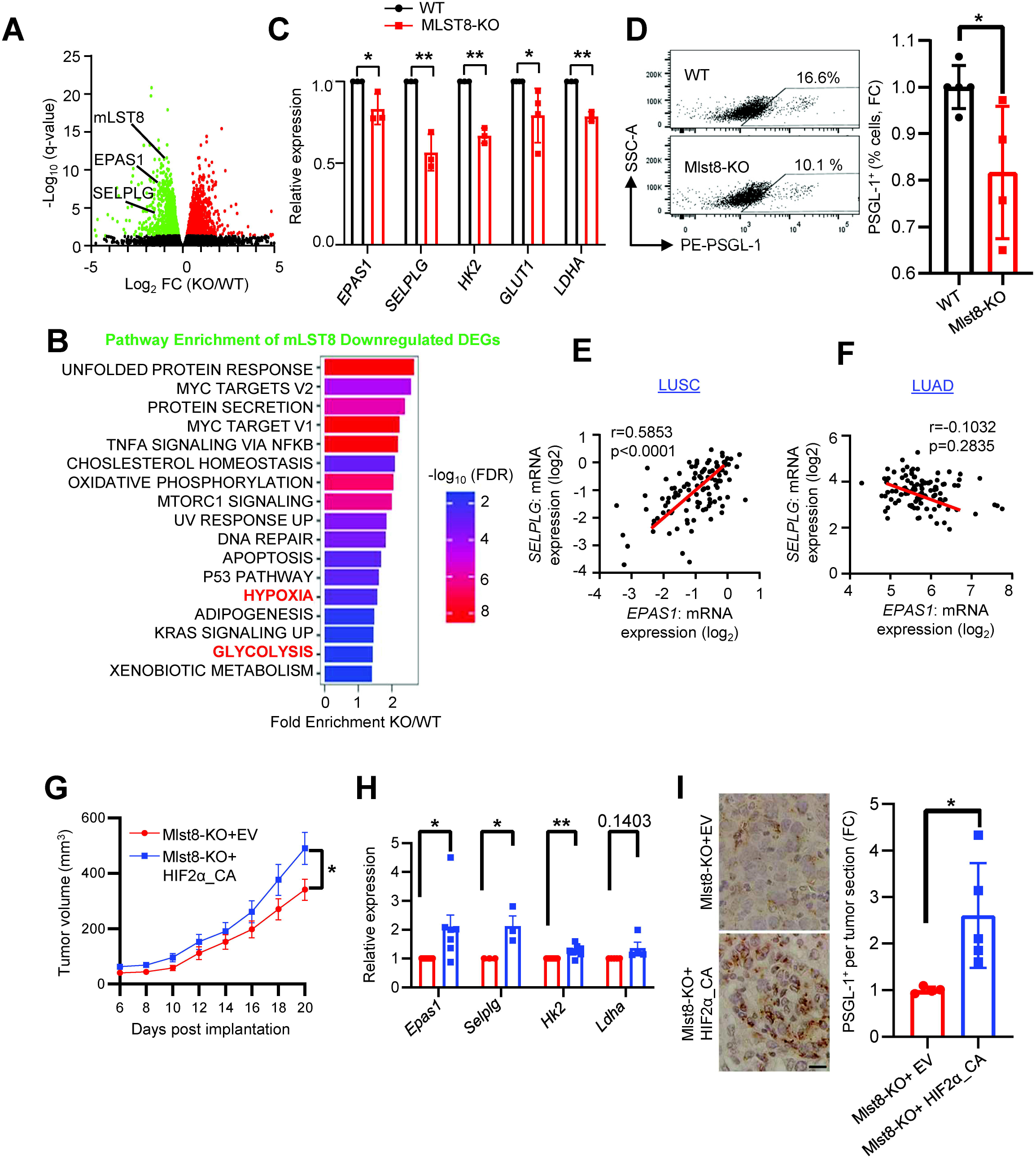
mTORC2 regulates PSGL-1/SELPLG via HIF2α/EPAS1. (A) Volcano plot showing differentially expressed genes from HCC2814 WT vs MLST8-KO cells (green=downregulated and red=upregulated). (B) Hallmark pathways presented for top 20 results based on fold enrichment, with an FDR cutoff of *p*<0.05 performed using the ShinyGO 0.77 bioinformatic tool. (C) mRNA expression of the indicated genes was validated by real-time PCR. (D) Flow cytometric analysis of PSGL-1^+^ (% cells, FC) from KLN205-Pten^null^ WT and Mlst8-KO cells in vitro. (E-F) Correlation between *SELPLG* and *EPAS1* using the LUSC CPTAC (E) and LUAD CPTAC datasets (F) downloaded from cBioPortal. Linear regression lines of best fit and Pearson’s correlation analyses are shown. (G) 5x10^5^ KLN205-Pten^null^ Mlst8-KO+EV or Mlst8-KO+HIF2α-CA cells were implanted in DBA/2 mice. Tumor growth was monitored, and tumor volume calculated (n=23 mice per group). (H) mRNA expression of the indicated genes validated by real-time PCR in KLN205-Pten^null^ Mlst8-KO+EV or Mlst8-KO+HIF2α-CA cells. (I) Immunohistochemistry and quantification of PSGL-1 expression on KLN205-Pten^null^ Mlst8-KO+EV or Mlst8-KO+HIF2α-CA tumor samples (n=4-5 mice per group). Scale bar: 20 μm. p-values were determined by the mixed-effects model (REML) with Sidak multiple comparisons post hoc (G) and by two-tailed unpaired Student *t* test (C, D, H, and I). *p<0.05, **p<0.01.

To investigate how mTORC2 may regulate *SELPLG*/PSGL-1 expression in tumor cells, we found that MLST8-KO cells exhibited a significant decrease in expression of *EPAS1*, the gene encoding HIF2α (Fig. 5A-C) which was previously shown to be regulated by mTORC2 (16,36). Indeed, in LUSC human patient samples, there is a strong positive correlation between *EPAS1*/HIF2α and *SELPLG*/PSGL-1 expression, but this association is lost in LUAD (Fig. 5E, F), indicating that the relationship between HIF2α and PSGL-1 may be specifically attributed to mTORC2 hyperactivation. To test if HIF2α rescues the MLST8-deficient phenotype in LUSC, we expressed a constitutively active mutant of HIF2α (HIF2α_CA) that is resistant to proteosome degradation in KLN205-Pten^null^ Mlst8-KO cells (Fig. S5E). Expression of HIF2α_CA accelerated tumor growth both in the flank and following orthotopic inoculation via intravenous injection (Figs. 5G, S5F-H). In addition, we found that HIF2α_CA rescued expression of the glycolysis genes *Hk2* and *Selplg*/PSGL-1 (Fig. 5H, I), suggesting that HIF2α may be a major transcriptional regulator downstream of mTORC2 in regulating glycolysis and PSGL-1 expression in LUSC.

### Anti-VISTA immunotherapy inhibits tumor growth and enhances antitumor immunity

Due to the absence of existing targeted therapies for mTORC2 and the modest success of immunotherapy in NSCLC (37–39), we sought to test if anti-VISTA immunotherapy can boost antitumor T cell activity and restrict LUSC growth. We orthotopically inoculated mice with KLN205-Pten^null^ WT or Mlst8-KO cells into the lungs of syngeneic mice via tail vein injection. After seeding, tumor-bearing mice were treated with an anti-VISTA antibody or IgG control (Fig. 6A). In animals treated with anti-VISTA, we observed a significant reduction in lung weights reflecting less tumor burden compared to those treated with IgG control (Fig. 6B). However, anti-VISTA does not affect tumor outgrowth in Mlst8-KO tumors (Fig. S6A, B), validating that VISTA and PSGL-1 function downstream of mTORC2. The reduction in tumor burden was associated with greater cleaved caspase3^+^ cells and an increase in CD8^+^ T cells in WT tumors without affecting Mlst8-KO tumors (Figs. 6C, S6C).

**Figure 6:**
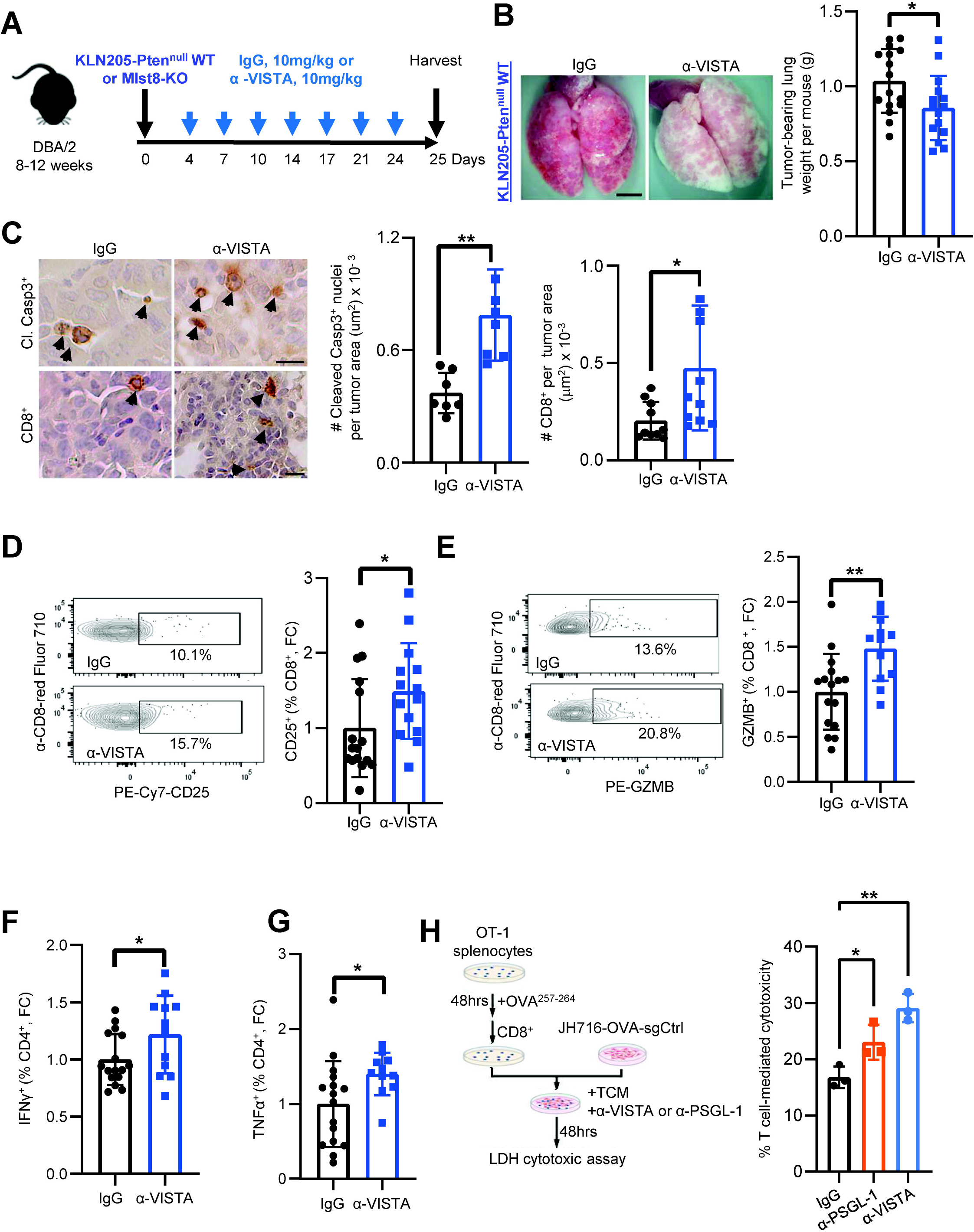
Systemic administration of anti-VISTA reduces lung tumor burden and promotes anti-tumor immunity. (**A**) Schematic of KLN205-Pten^null^ WT or Mlst8-KO cells inoculation and treatment schedule with IgG control and anti-VISTA (seven doses, blue arrows). (**B)** Representative images of the lungs harvested from IgG control or anti-VISTA treated KLN205-Pten^null^ WT tumor bearing mice after 25 days of tumor cell implantation. Scale bar: 100 μm. Lung weight indicating tumor burden from IgG Control and anti-VISTA. (**C**) Immunohistochemistry showing increased tumor cell apoptosis (Cleaved Caspase3) and increased CD8^+^ T cells in anti-VISTA treated animals. Scale bar: 20 μm. (**D-E**) Flow cytometry analysis of CD25^+^ (**D**), GZMB^+^ (**E**) on CD8^+^ from anti-VISTA treated mice. (**F-G)** Flow cytometry was further performed on IgG and anti-VISTA treated animals for IFNγ^+^ and TNF-α^+^ on CD4^+^ T cells. Each dot on the quantification represents one mouse. p-values were determined by two-tailed unpaired Student *t* test. (**H)** Schematic of cytotoxic co-culture assay and quantification, where JH716-OVA-sgControl cells were co-cultured with CD8^+^ T cells in tumor conditioned medium (TCM) and in the presence of IgG control, or α-VISTA, or α-PSGL-1 (n=3). p-values were determined by two-way ANOVA with Tukey’s post hoc. *p<0.05, **p<0.01, ****p<0.001, FC: Fold change.

Anti-VISTA treated tumors induced an increase in CD8^+^ T cell activation markers CD25^+^ and granzyme B (GZMB^+^) (Fig. 6D, E). Although we did not observe an increased in the number of CD4 T cells, there is an increased in CD4^+^IFN-γ^+^ and CD4^+^TNF-α^+^ cells, indicative of Th1 differentiation in the anti-VISTA treated group (Figs. 6F, G; S6D, E). Interestingly, there is also an increased in NK1.1^+^ cells and GZMB^+^NK1.1^+^ cells in α-VISTA treatment compared to IgG control (Fig. S6F, G). However, flow cytometric analyses revealed no changes in myeloid cells or macrophages, the T cell exhaustion marker TIM3, or PSGL-1 on tumor cells (Fig. S6H-L). Collectively, these data suggest that anti-VISTA enhances the effector function of tumor-infiltrating T cells, possibly by blocking the interaction between PSGL-1 and VISTA within the tumor microenvironment.

To test if anti-VISTA affects direct killing of tumor cells by T cells, we evaluated cytotoxicity of CD8+ T cells on the JH716-OVA cell line in the presence of VISTA or PSGL-1 neutralizing antibodies (Fig. 6H). The addition of either anti-VISTA or anti-PSGL-1 significantly increased tumor cell killing, compared to the IgG control group (Fig. 6H), suggesting a direct pro-cytolytic interaction of PSGL-1 on tumor cells or with VISTA on OT-I T cells. Taken together, these data suggest that targeting the immune checkpoint receptor VISTA or its ligand PSGL-1 may be an effective immunotherapy option for LUSC.

## Discussion

Targeted therapy has made a major impact in improving lung adenocarcinoma (LUAD) survival, but such success has not translated to lung squamous cell carcinoma (LUSC). One notable difference between LUAD and LUSC tumors is observed in metabolic dependencies. While LUAD tumors more heavily depend on oxidative phosphorylation (33), LUSC cells display elevated glucose metabolism (33,34). Targeting mTORC2 signaling by deletion of MLST8 reduced glycolysis and limited extracellular lactate levels of LUSC tumor cells both in vitro and in vivo, while also reducing tumor growth in mice. Activation of mTORC2 occurs downstream of PI3K signaling, and we show that genes in this signaling axis (*PIK3CA, PTEN, RICTOR*, or *AKT1-3*) are genetically altered in over 80% of LUSC patient samples. Therefore, this study supports mTORC2 as a new target for therapeutic intervention in LUSC.

While our results suggest the central role of mTORC2 in LUSC resides in regulation of glucose metabolism, mTORC2 has been found to control additional metabolic pathways (40,41). Although the role of mTORC2 in mitochondrial respiration is somewhat controversial (31), we observed a modest decrease in oxygen consumption rate in HCC2814 mLST8 KO cells. However, our results are consistent with the finding that mTORC2 is activated to maintain mitochondrial respiration, particularly under nutrient-limiting conditions (42,43). While the apparent multi-faceted role of mTORC2 in regulating metabolism may contribute to the LUSC phenotype, it is important to note that in vitro tumor cell viability and lymphocyte-deficient xenograft tumor growth was only marginally affected by mTORC2 status, suggesting that the intrinsic impacts of mTORC2 on tumor cell metabolism likely do not fully represent the complete landscape in LUSC. Indeed, our data indicates that mTORC2 in tumor cells has extrinsic influences on anti-tumor immunity in controlling LUSC tumor growth.

LUSC is strongly associated with smoking and a high mutation rate, favoring generation of neoantigens that can be recognized by cytotoxic T cells (44). Immune checkpoint blockade has been approved for NSCLC, but response rates in LUSC have not been optimal. The median survival of LUSC patients treated with a combination of anti-PD-1 and/or anti-PD-L1 with chemotherapy is only 17.1 months (37–39,45), necessitating the need to improve immunotherapy options. Based on our discovery, targeting mTORC2 is expected to inhibit glycolysis and restore a more neutral pH in tumor microenvironment. This will have major benefits for antitumor T cells in at least two respects. Lactate has been shown to serve as an oncometabolite, suppressing T cell motility, inhibiting T cell cytotoxicity and effector function, and promoting M2 macrophage differentiation (46–48). In addition, an acidic pH facilitates the binding of VISTA to PSGL-1, further driving immune suppression (7). Interestingly, loss of MLST8/mTORC2 signaling also decreases the expression of PSGL-1 on tumor cells, serving to further influence T cell activity through VISTA. Indeed, treatment with an anti-VISTA antibody significantly reduced LUSC tumor growth in WT, but not mLST8 KO cells, indicating that both the mTORC2-dependent effects, including those that drive glycolysis and PSGL-1 expression have dramatic impacts on LUSC, at least in part, through modulating anti-tumor immune response.

Unfortunately, there are currently no FDA approved small-molecule inhibitors that selectively block mTORC2 activity. Dual inhibitors that inhibit the mTOR kinase in both mTORC1 and mTORC2 have been proven to be poorly suitable due to the immunosuppressive nature of mTORC1 inhibition in immune cells (49,50). In addition, mTOR kinase inhibitors release negative feedback on PI3K/Akt in tumor cells (51). However, several new strategies for selective inhibition of mTORC2 offer great promise for enhancing immunotherapy in LUSC. Nanoparticles carrying siRNA against Rictor, a unique component of mTORC2, were shown to inhibit breast cancer tumor growth in vivo (52). It will be important to test whether these nanoparticles against mTORC2 are also effective in treatment of LUSC, either as a single agent or in combination with existing immunotherapies. Furthermore, we have shown that LUSC is sensitive to anti-VISTA treatment in an orthotopic LUSC animal model. VISTA is expressed in most LUSC tumors, with levels correlating to increased lymphocyte infiltration and PD-1 expression (53). This co-expression of VISTA and PD-1 may further support poor anti-tumor T cell responses given the apparent non-redundant role of these two checkpoint receptors. Therefore, there is great potential that combining anti-VISTA with anti-PD-1 may improve current responses of LUSC patients that are not responsive to first- or second-line immunotherapy. Finally, the HIF2α inhibitor Belzumtifan is FDA approved for treating renal cell carcinoma (54). Because HIF2α plays an important role to regulate the expression of *SELPLG*/PSGL-1 and glycolytic enzymes downstream of mTORC2, it will be interesting to test if this inhibitor is also effective in inhibiting LUSC tumor growth and whether it can improve the current immunotherapy.

## Acknowledgements

We thank Dr. Christine Lovely (Vanderbilt University Medical Center) for providing clinical expertise and helpful discussion and Dr. Chad Pecot (University of North Carolina, Chapel Hill, NC) for providing the JH716 cells. This work is supported by VA Career Scientist Award 5IK6BX005391 and a VA Merit Award 5101BX000134 (to J.C.). J.C. is also supported by NCI grants CA250506 and CA271176. D.N.E. is supported by a Department of Defense CDMRP W81XWH2210109 and Dr. Verra Ngwa is supported by an AACR/Johnson & Johnson postdoctoral fellowship. Flow cytometry experiments were performed in the Vanderbilt University Medical Center (VUMC) Flow Cytometry Shared Resource, which is supported by the Vanderbilt Ingram Cancer Center (P30 CA68485) and the Vanderbilt Digestive Disease Research Center (DK058404). Tissue processing and H&E staining was performed by the VUMC Translational Pathology Shared Resource, which is supported by NCI/NIH Cancer Center Support Grant P30CA068485.

## Contributions

V.M.N, Y.H., and J.C. conceptualized the project and developed methodologies. V.M.N, Y.H., D.N.E., and W.S. performed experiments. Data analysis and interpretation was performed by V.M.N., Y.H., D.N.E., and J.C. V.M.N, D.N.E, and J.C. wrote, reviewed, and/or revised the manuscript.

