## Supplementary material for "Targeting mTORC2 in lung squamous cell carcinoma improves anti-tumor immunity through the PSGL-1-VISTA axis": Supplemantal _.pdf

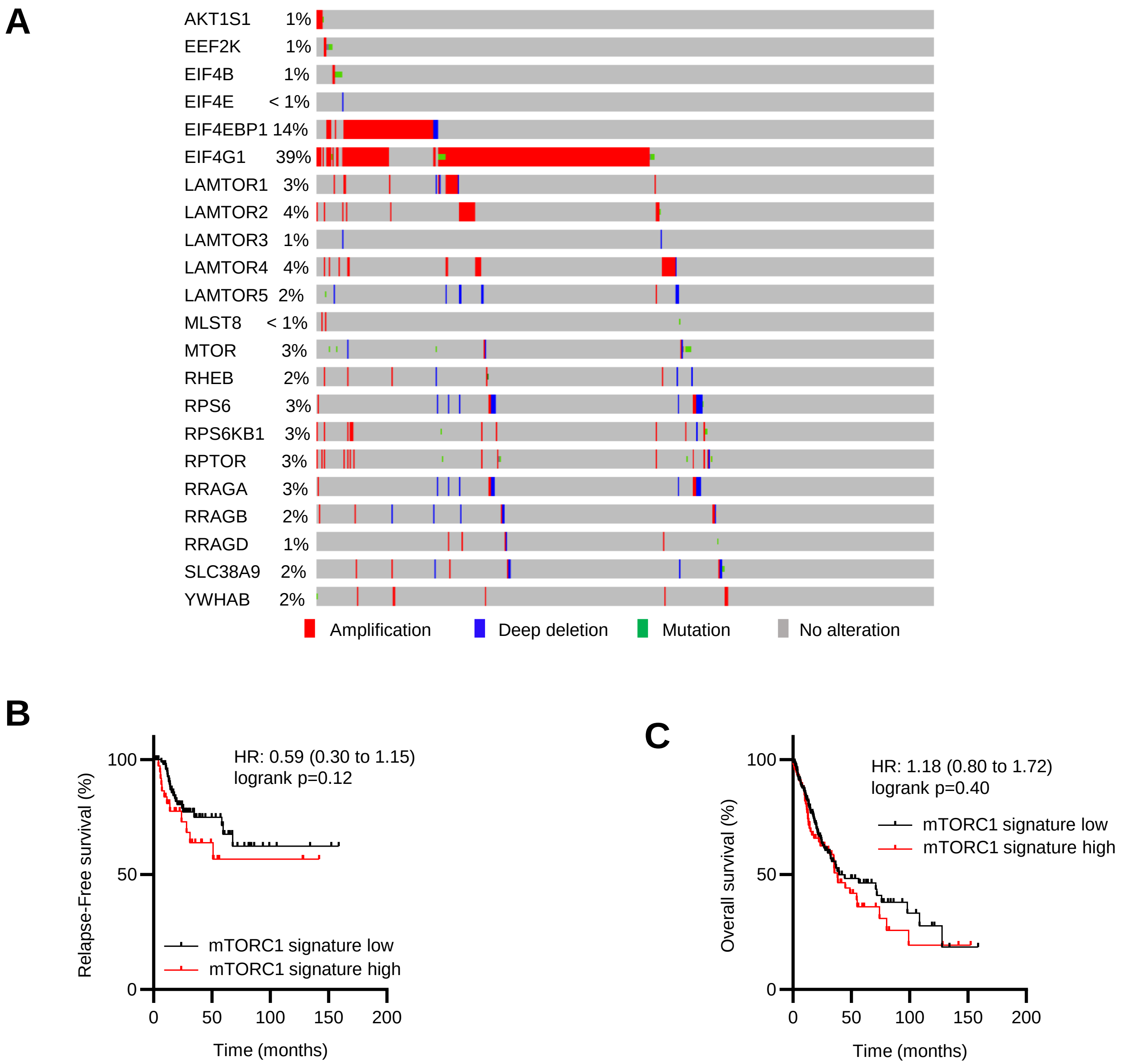
A
AKT1S1 1%
EEF2K 1%
EIF4B 1%
EIF4E < 1%
EIF4EBP1 14%
EIF4G1 39%
LAMTOR1 3%
LAMTOR2 4%
LAMTOR3 1%
LAMTOR4 4%
LAMTOR5 2%
MLST8 < 1%
MTOR 3%
RHEB 2%
RPS6 3%
RPS6KB1 3%
RPTOR 3%
RRAGA 3%
RRAGB 2%
RRAGD 1%
SLC38A9 2%
YWHAB 2%
Amplification
Deep deletion
Mutation
No alteration
B
HR: 0.59 (0.30 to 1.15)
logrank p=0.12
Relapse-Free survival (%)
mTORC1 signature low
mTORC1 signature high
Time (months)
C
HR: 1.18 (0.80 to 1.72)
logrank p=0.40
Overall survival (%)
mTORC1 signature low
mTORC1 signature high
Time (months)

**D**

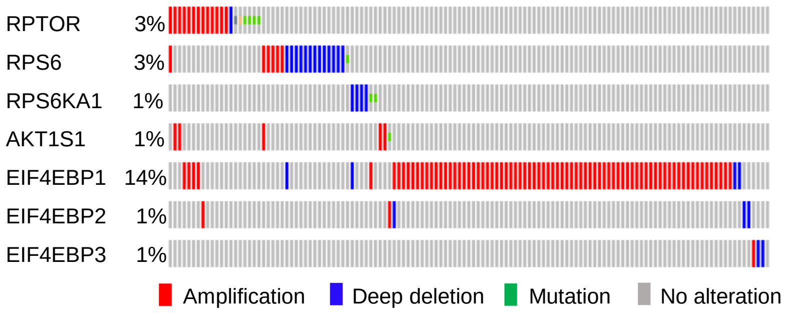


**E**

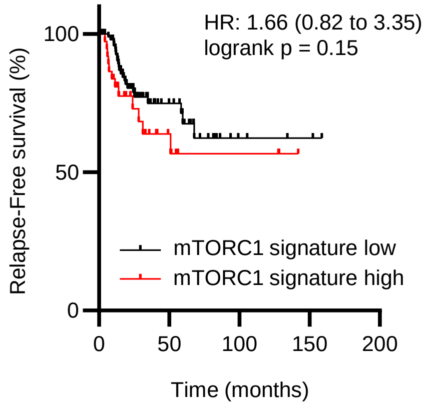


**F**

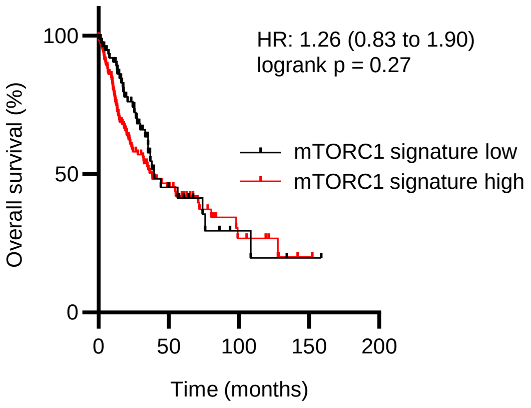


**Supplementary Figure 1 related to Figure 1: Genetic alterations in LUSC data analyzing mTORC1 genes.** (**A**) Analysis of mTORC1 related genes in the REACTOME_MTORC1_MEDIATED_SIGNALLING gene set in LUSC datasets [**LUSC:** CPTAC (n=80), TCGA Firehose Legacy (n=511)] using cBioPortal. (**B-C**) Kaplan-Meier (KM) of stage II-IV plot displaying the probability of relapse-free survival (**B**) and overall survival (**C**) with association of the mTORC1 related genes illustrated in **A** using lung squamous cell carcinoma RNA-seq datasets (n=141, relapse-free survival; n=249, overall survival) downloaded from kmplot.com. (**D-F**) mTORC1 unique components and downstream genes (mTORC1 signature: *RPTOR*, *RPS6*, *RPS6K*, *A1AKT1S1*, *EIF4EBP1*, *EIF4EBP2*, *EIF4EBP3*) were further analyzed using the same datasets as above. Log-rank (Mantel-Cox) test *p*-values, Hazard ratio (HR) (log-rank) and 95% confidence intervals are shown.

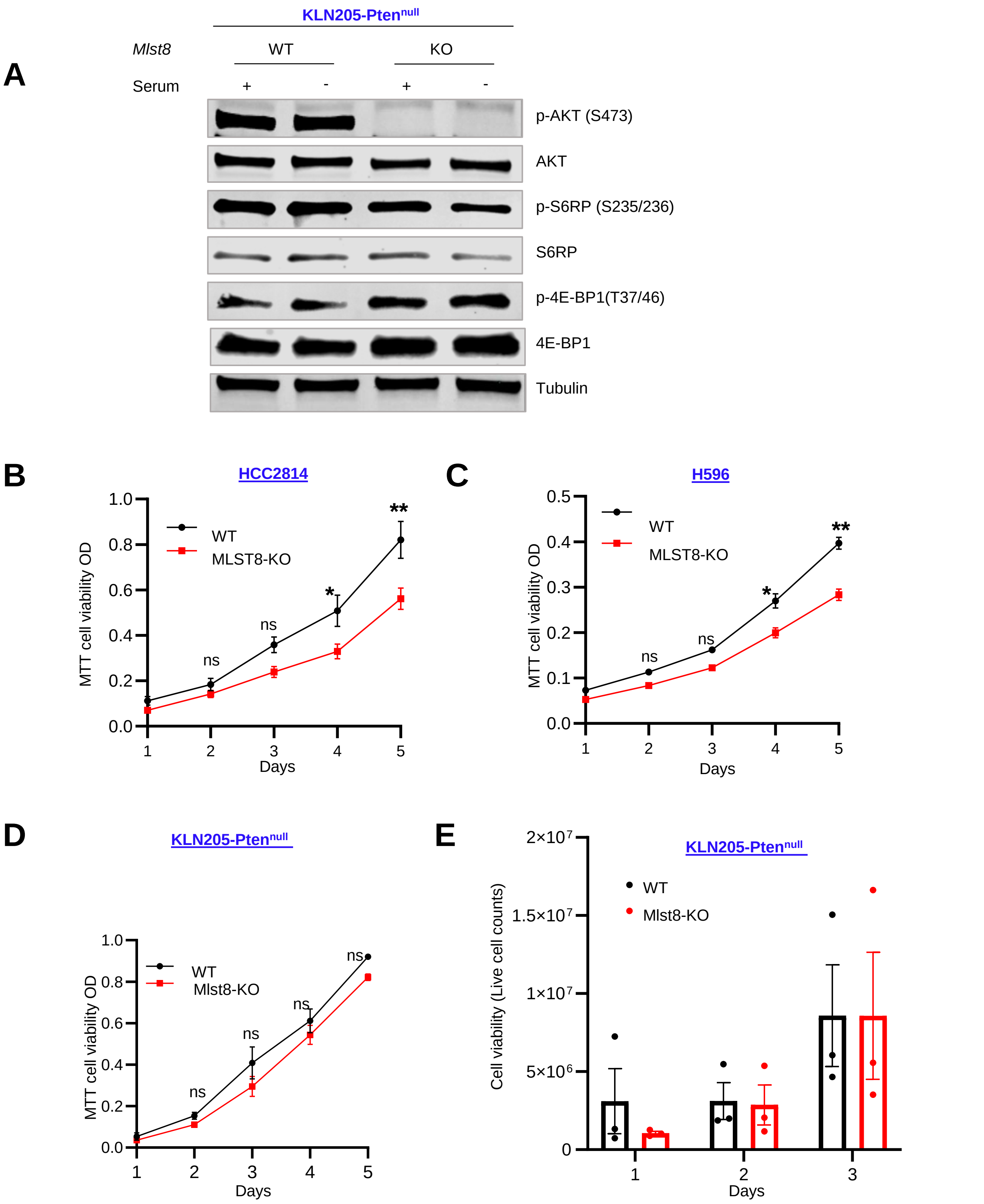


**Supplementary Figure 2 related to Figure 2: LUSC invitro cell proliferation**. (**A**) KLN205-Pten$^{null}$ WT or Mlst8-KO cells were serum starved overnight, and cell lysates were assessed by western blot analysis. (**B-C**) Cell viability was measured by MTT assay in HCC2814 and H596 WT or MLST8-KO cells (n=3 biological replicates). (**D**) KLN205-Pten$^{null}$ WT or Mlst8-KO cells were cultured for a total of 72 hrs, and dead cells were stained with trypan blue. All data are presented as mean ± SEM from two or three independent experiments. p-values were determined by 2-way ANOVA with Sidak multiple comparisons correction test. *p<0.05, **p<0.01, ns: not statistically significant.

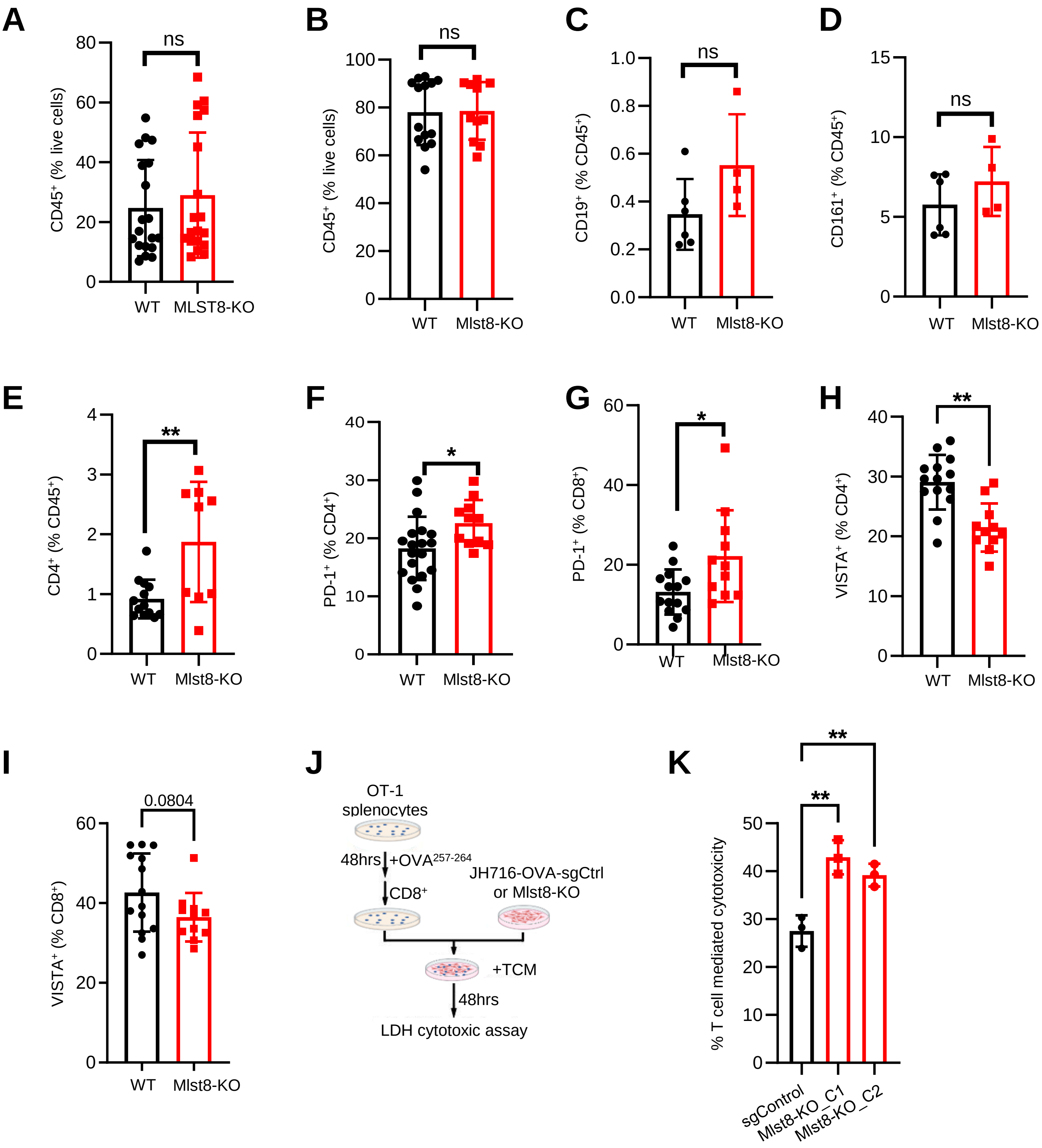

A
CD45⁺ (% live cells)
ns
WT
MLST8-KO
B
CD45⁺ (% live cells)
ns
WT
Mlst8-KO
C
CD19⁺ (% CD45⁺)
ns
WT
Mlst8-KO
D
CD161⁺ (% CD45⁺)
ns
WT
Mlst8-KO
E
CD4⁺ (% CD45⁺)
**
WT
Mlst8-KO
F
PD-1⁺ (% CD4⁺)
*
WT
Mlst8-KO
G
PD-1⁺ (% CD8⁺)
*
WT
Mlst8-KO
H
VISTA⁺ (% CD4⁺)
**
WT
Mlst8-KO
I
VISTA⁺ (% CD8⁺)
0.0804
WT
Mlst8-KO
J
OT-1 splenocytes
48hrs
+OVA257-264
CD8⁺
JH716-OVA-sgCtrl or Mlst8-KO
+TCM
48hrs
LDH cytotoxic assay
K
% T cell mediated cytotoxicity
**
**
sgControl
Mlst8-KO_C1
Mlst8-KO_C2

**Supplementary Figure 3 related to Figure 3: Effect of MLST8/mTORC2 loss on immune landscape. (A-D**) Flow cytometry quantification of $CD45^+$ in H596 tumors (**A**), $CD45^+$ (**B**), B- and NK cells (**C, D**) in KLN205-$Pten^{null}$ WT or Mlst8-KO tumors (n=4-14). (**E**) Flow cytometric analysis of $CD4^+$ (% $CD45^+$) from WT and Mlst8-KO tumors. (**F-I**) Flow cytometric analysis of $PD1^+$ (% $CD4^+$), $PD1^+$ (% $CD8^+$), $VISTA^+$ (% $CD4^+$), $VISTA^+$ (% $CD8^+$) from WT and Mlst8-KO tumors. Each dot represents a mouse. (**J-K**) Cytotoxicity of HJ716-OVA cells was determined. (**J**) Schematic of the co-culture assay is shown. (**K**) JH716-OVA-sgControl vs Mlst8-KO (C1: clone_1 and C2: clone_2) cells were co-cultured with $CD8^+$ T cells in tumor conditioned medium (TCM) (n=3). All data are presented as mean ± SEM from two or three independent experiments. p-values were determined by two-tailed unpaired Student *t* test. *$p<0.05$, **$p<0.01$. ns: not statistically significant.

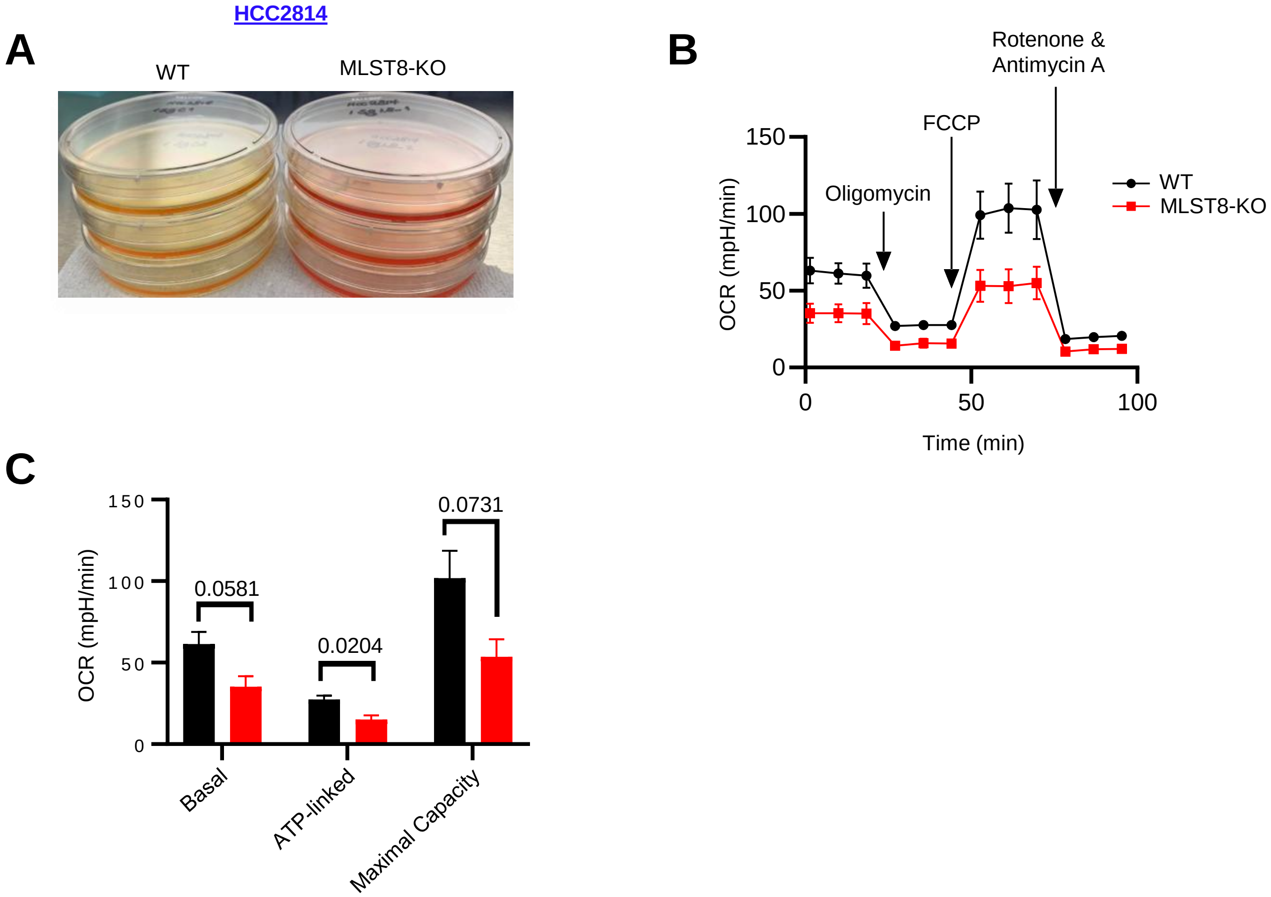


**Supplemental Figure 4 related to Figure 4: Loss of function in mTORC2 alters mitochondrial respiration.** (**A**) Representative image of media color changes in HCC2814 WT and MLST8-KO cell. (**B**) Oxygen Consumption Rate (OCR) changes in HCC2814 WT and MLST8-KO cells after mLST8 was deleted. (**C**) Bar chart showing mitochondrial respiration function parameters of WT versus MLST8-KO HCC2814 cells analyzed with basal respiration, ATP production and maximal respiration. All data are presented as mean ± SEM from two or three independent experiments. p-values were determined by two-tailed unpaired Student *t* test.

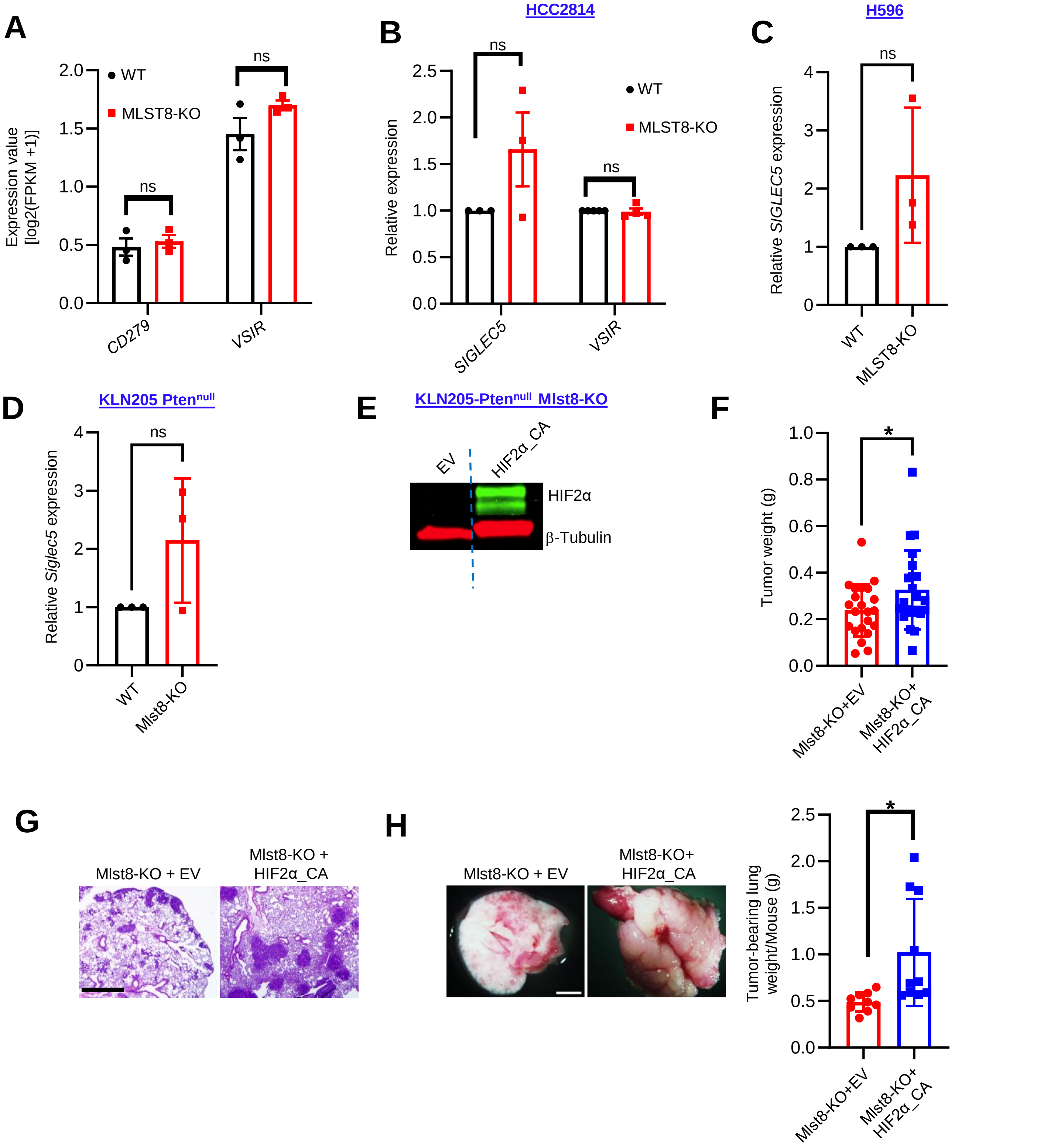


**Supplementary Figure 5 related to Figure 5: mTORC2 loss of function does not change other PSGL-1 binding partners but regulates PSGL-1 expression through HIF2α**. (**A**) Expression values of *PD-L1* and *VISTA* in MLST8-KO compared to WT from RNA-sequencing. (**B-D**) Expression values of *VISTA* and *SIGLEC-5* by RT-qPCR from HCC2814, H596 and KLN205-Pten$^{null}$ WT or MLST8-KO cells (n=3). (**E**) Western blot of HIF2α from KLN205-Pten$^{null}$ Mlst8-KO cells transfected with Empty Vector (EV) or HIF2α-CA. Blue dotted line denotes spliced gel. (**F**) Subcutaneous tumor weight. (**G-H**) 1x10$^{6}$ KLN205-Pten$^{null}$ Mlst8-KO+ EV or Mlst8-KO+ HIF2α-CA cells were injected into DBA/2 mice via tail vein. Representative H&E staining (**G**) and whole lung and weight (**H**) harvested from Mlst8-KO+EV or Mlst8-KO+HIF2α_CA tumor bearing mice after 21 days. Scale bar: 100 μm. Each dot represents a mouse. All data are presented as mean ± SEM from two or three independent experiments. p-values were determined by two-tailed unpaired Student *t* test. *p<0.05, ns: not statistically significant.

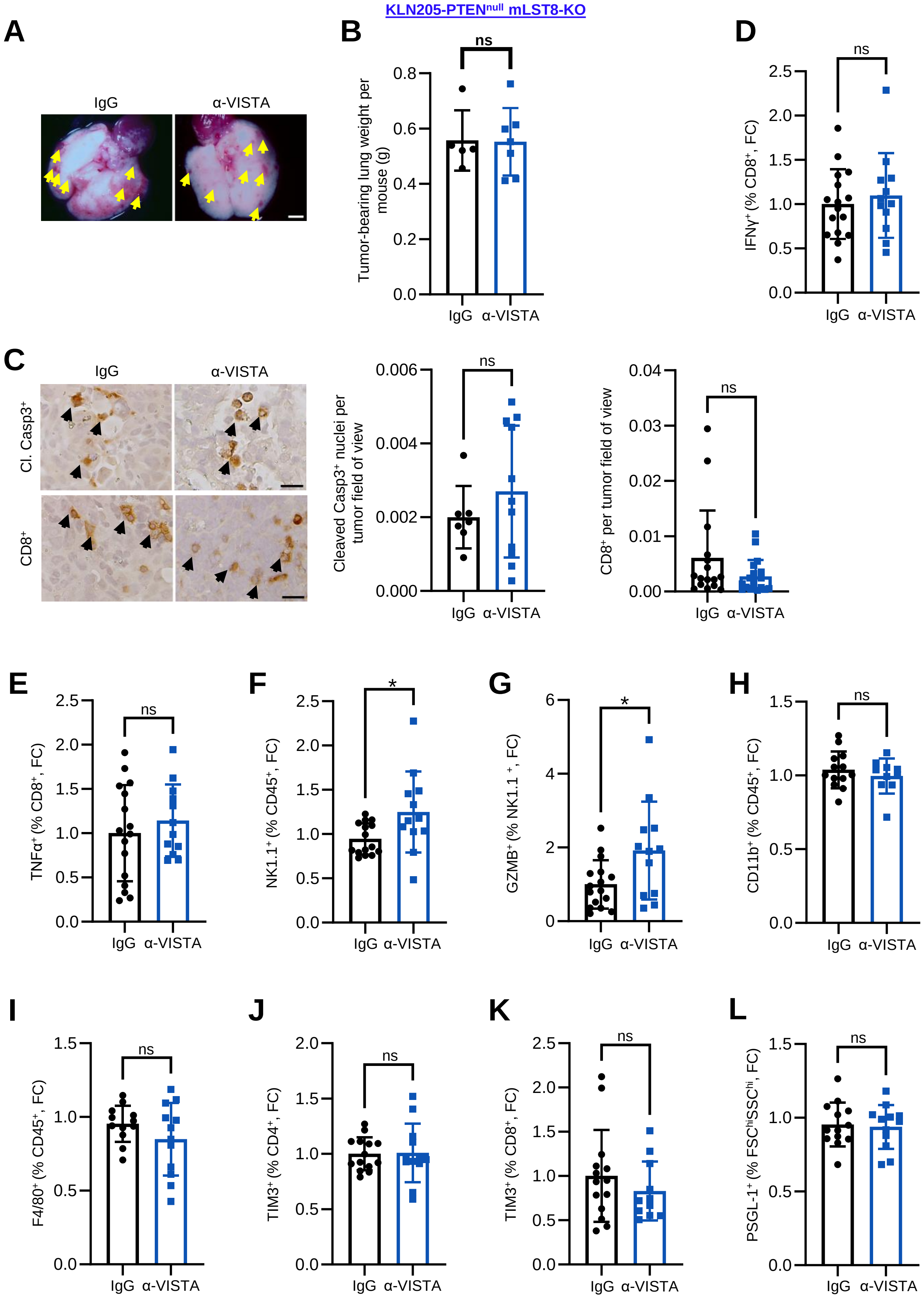

KLN205-PTEN^null mLST8-KO
A
IgG
α-VISTA
B
Tumor-bearing lung weight per mouse (g)
ns
IgG α-VISTA
D
IFNγ+ (% CD8+, FC)
ns
C
Cl. Casp3+
CD8+
Cleaved Casp3+ nuclei per tumor field of view
CD8+ per tumor field of view
E
TNFα+ (% CD8+, FC)
F
NK1.1+ (% CD45+, FC)
*
G
GZMB+ (% NK1.1+, FC)
H
CD11b+ (% CD45+, FC)
I
F4/80+ (% CD45+, FC)
J
TIM3+ (% CD4+, FC)
K
TIM3+ (% CD8+, FC)
L
PSGL-1+ (% FSC^hiSSC^hi, FC)

**Supplementary Figure 6 related to Figure 6: Anti-VISTA treatment in vivo and in vitro improves T cell cytotoxic killing.** (**A**) Representative images of the lungs harvested from IgG control or anti-VISTA treated KLN205-$Pten^{null}$ Mlst8-KO tumor bearing mice after 25 days of tumor cell implantation. Scale bar: 100 µm. (**B**) Lung weight indicating tumor burden from IgG Control and anti-VISTA. Each dot on the quantification represents a mouse. (**C**) Immunohistochemistry identifying Cl. Caspase $3^+$ and $CD8^+$ cells on IgG vs anti-VISTA tumor samples. Scale bar: 50 µm. Each dot on the quantification represents Cl. Caspase $3^+$ or $CD8^+$ nuclei per tumor field of view. (n=4-6 mice per group). (**D-L**) Flow cytometry was performed on $IFN\gamma^+CD8^+$ (**D**), TNF-$\alpha^+CD8^+$ (**E**), $NK1.1^+CD45^+$ natural killer cells (**F**), $GZMB^+NK1.1^+$ (**G**), $CD11b^+$ $CD45^+$ myeloid cells (**H**), $F4/80^+CD45^+$ macrophages (**I**), $TIM3^+CD4^+$ (**J**), $TIM3^+CD8^+$ (**K**), and PSGL-$1^+$ (**L**) from IgG vs anti-VISTA treated mice. Each dot on the quantification represents a mouse. p-values were determined by two-tailed unpaired Student *t* test. *$p<0.05$, ns: not statistically significant.
